# Diverging effects of global change on future invasion risks of *Agama picticauda* between invaded regions: same problem, different solutions

**DOI:** 10.1101/2025.08.21.671467

**Authors:** Nicolas Dubos, Steven Calesse, Kathleen C. Webster, Chloé Bernet, Gregory Deso, Thomas W. Fieldsend, Saoudati Maoulida, Mounir Soulé, Xavier Porcel, Jean-Michel Probst, Hindatou Saidou, Jérémie Souchet, Mohamed Thani Ibouroi, Markus A. Roesch

**Author notes:** Corresponding author: Nicolas Dubos. equal contribution (Mohamed Thani Ibouroi, Markus A. Roesch).

## Abstract

Predicting biological invasions is challenging because multiple factors can act in contrasting directions and exert heterogeneous effects across space. Nevertheless, modelling approaches provide valuable tools to anticipate the potential spread of invasive alien species and to support mitigation strategies. With an Ecological Niche Modelling approach, we predicted the invasion risks of Peters’s Rock Agama *Agama picticauda*, a species that is spreading globally in non-forested areas through freight transport and un-/intentional releases from the pet trade. The potential establishment of the species in new areas is of concern for multiple endemic species throughout the world. We quantified the effects of climate, anthropogenic activity and forest cover on invasion risk. We used verified records from the native and non-native range and accounted for the latest methodological recommendations. We predicted how invasion risk will vary in the future (2070) using projections from two scenarios (SSP2 and SSP5). We predict that invasion risks will vary in diverging directions, depending on the region. The risk will increase in human-populated regions and on small islands but will decrease in Florida. We recommend increasing surveillance in vehicular transportation of material especially within the Comoros and the Mascarenes archipelagos. Since many introductions are related to the pet trade in Florida, we recommend stronger legal regulations and the promotion of public awareness. Promoting tree cover may be locally beneficial to prevent establishment of *A. picticauda*. The effect of climate change, land use change and human activities may differ between and within both, the native and the invaded regions.

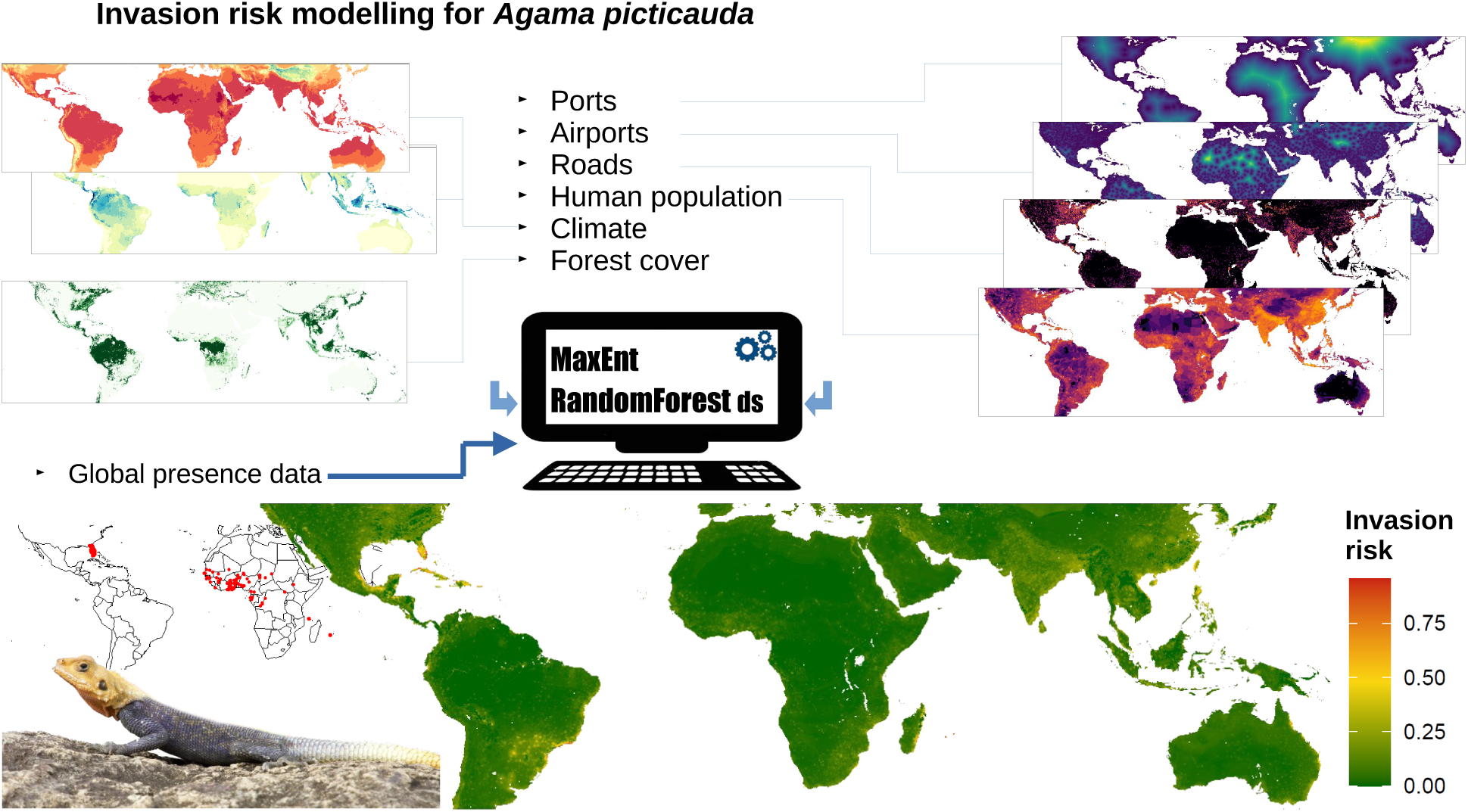

## 1. Introduction

Global transportation of goods as well as the international pet trade have led to many incidental or deliberate introductions of non-native species throughout the world (Gippet and Bertelsmeier, 2021). Once introduced to new environments, some species can establish and colonize further grounds, causing severe ecological and economic impact (Bellard et al., 2016; Diagne et al., 2021). New introductions are being reported every year, with no sign of saturation whatsoever (Clements et al., 2025; Seebens et al., 2017). Invasive alien species (IAS) are one of the main causes of native species decline, and the risk of establishment and spread of IAS is likely to vary in the future with global change (Bellard et al., 2013). Climate change can affect invasion risks in either direction, i.e., increasing or reducing the risk (Bellard et al., 2013; Dubos et al., 2023a), and the effect of climate and anthropogenic activities may sometimes also be contrasting (e.g., Dubos et al., 2025). Land use change can also alter invasion risk either positively or negatively, depending on the ability of the species to exploit anthropogenic structures and resources (e.g. Mitchell et al., 2021). Environmental change can affect populations of the native and the non-native region in opposite directions, due to different socio-economic contexts (Dubos et al., 2025). Similarly, for IAS that have spread globally, the mechanisms of invasion may differ between the invaded regions. However, few studies report quantifications of invasion risk while distinguishing between invaded regions, which can make management recommendations unsuitable to the local context in some cases.

The best strategy to mitigate the ecological and economic impacts caused by IAS is the prevention of new introductions (Pearson, 2024). This can be achieved with the promotion of proactive surveillance and early detection at the main entry points such as airports and harbours in target regions (Cuthbert et al., 2022). The regions that are the most at risk in the present and future can be identified with predictive modelling approaches such as Ecological Niche Models (ENMs; also known as Species Distribution Models, SDMs). These models are widely used to predict species response to future environmental change (e.g., Dubos et al., 2023b; Rose et al., 2024; Serva et al., 2023; Wiens and Zelinka, 2024), prioritizing conservation areas (e.g., Domisch et al., 2019; Dubos et al., 2022b; Faleiro et al., 2013; Giakoumi et al., 2025; Leroy et al., 2014), testing ecological hypotheses (Raxworthy et al., 2007) or assessing invasion risk (Gallien et al., 2012). However, increasing accessibility with user-friendly interfaces of highly complex models designs has led to a widespread misuse or misinterpretation of SDMs (e.g. ignorance of biological or statistical assumptions, subjective reliance on performance metrics, systematic use of default parameters; Leroy, 2023; Yackulic et al., 2013; Zurell et al., 2020). As a result, projections of future invasion risk can largely differ and become potentially misleading. However, provided the model design is in line with ecological theory and results are interpreted specific to the natural history of the study organism, SDMs can produce maps that help target priority entry points of potential IAS for stakeholders (Hui, 2022).

Modelling the environmental niche of non-native species that are still spreading is challenging because of the non-equilibrium hypothesis (Gallien et al., 2012; Hui, 2022). After being introduced outside of their native range, IAS have the potential to extend their realized niche as a result of ecological release (i.e. fewer biotic constraints) and in situ adaptation (Fieldsend et al., 2021; Refsnider et al., 2015). At an early stage of invasion, the species is not at equilibrium and is susceptible to extend further its range. Hence the available occurrence data does not fully represent the true extent of suitable environments that are likely to be colonized. Therefore, models might downplay the importance of some suitable regions that have not yet been invaded and underestimate the potential spread. To mitigate this effect, it is recommended to include both data from the native and non-native range to train the models with all available information on the ecological niche of the species (Barbet-Massin et al., 2018). It is further possible to exclude zones that have not yet been reached by the species from the model training to reduce model omission error and the effect of non-equilibrium (Chapman et al., 2019). To do so, the model training area (i.e. background) should be limited to a restricted area near the known occurrences, and more weight should be given to the most visited places (i.e. the equivalent of correcting for sample bias). Eventually, since the non-native species could be introduced into places that are spatially separated, it is of critical importance to evaluate models using approaches that enable the assessment of spatial transferability (e.g. block-cross validation). To maximize the reliability of predictions of invasion risks, models must be designed in close interaction between modellers and experts.

Peters’s rock agama *Agama picticauda*, also known as the African red-headed agama or African rainbow lizard, is native to central-western Africa (Leaché et al., 2014). Its current global distribution is the result of both incidental escapes from the pet trade, deliberate introductions from pet owners, hitchhiking with maritime transportation of goods and natural dispersion through natural or anthropogenic structures (Gray, 2020; Guillermet et al., 1998; Mitchell et al., 2021). The species was confounded with *A. agama* species complex until its systematic and taxonomic clarification by Leaché et al. (2014) The identity of introduced populations of this agama has also been confirmed as *A. picticauda* (Nuñez et al., 2016; Roesch et al., 2025). The species is now present and well-established outside of its native range in Florida, Réunion Island and Grand Comore, where it may represent a threat to local species. Further introductions have already begun to be documented in the Bahamas, the British Virgin Islands, Madagascar, Cape Verde Islands and the USA (where establishment is yet to be confirmed; Enge, 2024; Wagner et al., 2009 and references therein). Additional introductions and establishments are expected to occur in presently unreached regions (e.g. in the Lesser Antilles; van den Burg et al., 2024).

*Agama picticauda* can be found in a variety of habitat types in its native region, but inhabits mostly open habitats such as savannah or forest edges (Krishnan et al., 2019). The species particularly benefits from anthropogenic structures, where it can find shelter, thermoregulation sites, a high prey availability and few competitors (Mitchell et al., 2021). The species is an opportunistic insectivore, and introduced populations have recently been found to show dietary niche conservatism preserving Hymenoptera as their main prey (Harris et al., 2025). However, it has also been observed eating fruits or discarded human food items (Gray, 2020; Ofori et al., 2018). The preference for hymenopteran diet and the high flexibility of the species to exploit anthropogenic food sources grants it a concerning potential for further spread in the tropics.

The presence of *A. picticauda* could represent a major threat to native reptiles. There is evidence of a risk of predation on smaller individuals (including cannibalism and predation on reptiles; Henigan et al., 2019; Ofori et al., 2023; van den Burg et al., 2024). The species may also impact native reptiles through interspecific competition (van den Burg et al., 2024)*. Agama picticauda* displays an aggressive territoriality to which endemic reptiles can be highly sensitive (e.g. *Phelsuma inexpectata* in Réunion Island; Deso et al., 2023). Multiple regions are of concern, as for example in Grand Comore, where the endemic *Oplurus cuvieri comorensis* is already the focus of conservation efforts (Hawlitschek et al., 2011). On Réunion Island, *A. picticauda* has rapidly spread across all coastal areas since its introduction (first record in 1995; Guillermet et al., 1998) and is progressing up to higher elevations (Roesch et al., 2025). The positive effect of anthropogenic structures is even more worrying for species that dwell within urban areas such as the critically endangered *P. inexpectata*, or those living in the vicinity of villages such as *P. borbonica* (Augros et al., 2017; Dubos, 2013; Dubos et al., 2014). Its introduction is expected to affect many reptile species in the Lesser Antilles (> 45 species; van den Burg et al., 2024). The threat to endemic species and the ongoing spread of the species call for the urgent need to predict invasion risks at the global scale.

*Agama picticauda* is absent from dense forests, suggesting a lower impact on natural forested areas (van den Burg et al., 2024). However, the high transition rate of forests to urban areas (e.g. in the Caribbean; van den Burg et al., 2024) and the alarming rates of deforestation at the global scale suggest a potential increase in the area of habitat suitable for this species in the future. We present this study as a response to an urgent need to identify areas that are predicted to be suitable for *A. picticauda* by 2070, accounting for future projections of climate, land use and human activity.

We modelled the invasion risks of *A. picticauda*, accounting for its environmental niche (climate and habitat) and factors of introduction and spread. With a robust approach which accounts for the latest methodological recommendations, we identify regions that are the most at risk, quantify the effects of future climate change and deforestation for the native and the three invaded regions separately, and provide guidelines for targeted future surveillance efforts.

## 2. Methods

### 2.1 Overview

we used an ecological niche modelling approach with a combination of the two best performing algorithms (RandomForest down-sampled and MaxEnt; details and references below), correcting for sample bias (coping with non-equilibrium assumptions), with a spatial partitioning for model evaluation (to assess spatial transferability), a robust predictor selection process, accounting for anthropogenic factors, habitat and high-resolution global climate data that are appropriate for small islands.

### 2.2 Occurrence data

We used verified occurrence data from multiple sources from both the native and non-native regions, as recommended to best inform the models of the environmental conditions already filled by the species (Hui, 2022). We selected verified data points from multiple sources: i.e. the Global Assessment of Reptile Distributions (GARD; Roll et al., 2017), GBIF (gbif.org, <u>excluding preserved specimen and coordinates with a precision > 1km</u>), iNaturalist research grade, the association Nature Océan Indien (NOI), Système d’information de l’inventaire du patrimoine naturel (SNIP La Réunion; borbonica.re), and from the Early Detection & Distribution Mapping System (EDDMaps; eddmaps.org) and from published and unpublished literature (Augros et al., 2019; Leaché et al., 2017). For the native range, we only used data from Leaché et al. (2014, 2017) to avoid the inclusion of misidentified agamas. We complemented those data with further recent opportunistic field observations from Réunion Island and a dedicated survey in the Comoros. The latter was conducted across the entire island of Grande Comore between June and November 2024, where administrative regions were surveyed (Soule et al., 2025). We performed our sampling across a range of habitats to discriminate suitable and non-suitable habitats. In each location, presence data were gathered (when available) along trails, roads, or along line transects established both in intact and degraded natural forests. Whenever possible, we selected routes that crossed different habitat types (e.g., urban areas, agricultural land, or forest) to maximise the likelihood of detecting agama across a variety of environments within each geographical area.

This results in an initial sample size of unique point coordinates of 4140 points (native + non-native ranges). To avoid pseudo-replication, we resampled one point per pixel of environmental data at 1 km resolution (i.e. a process of spatial data thinning; (Steen et al., 2021), resulting in a final sample of 1289 points (90 from the native range and 1199 from the non-native range).

### 2.3 Climate data

Predictions of climate suitability, as well as invasion risks, can be highly sensitive to the choice of climate data source (Dubos et al., 2023a, 2022a). We used 19 bioclimatic variables (description available at https://www.worldclim.org/data/bioclim.html; see also Booth et al., 2014) at 30 arc seconds (approximately 1 × 1 km) resolution for the current (1979-2013) and future (2061-2080, hereafter 2070, i.e. the midpoint) climate from CHELSA version 1.2 (Karger et al., 2017; CMIP5, obtained at https://chelsa-climate.org). There is a significant uncertainty related to the source of climate data in ENM, and it is recommended to use multiple climate data sources (Baker et al., 2016; Dubos et al., 2022a). Worldclim is the only alternative available at the global scale in high resolution (1 × 1 km) and with projections of future conditions for multiple scenarios and Global Circulation Models, GCMs (Fick and Hijmans, 2017). However, Worldclim is known to be less reliable in small oceanic islands (IPCC, 2021), which is why we preferred to use CHELSA only, which better takes into account orographic factors.

For future projections, we used three commonly used GCMs (GCMs; BCC-CSM1-1, MIROC5, and HadGEM2-AO) to account for uncertainty in the methodology. We used two greenhouse gas emission scenarios (one optimistic RCP45-SSP2 and the most pessimistic one RCP85-SSP5) to consider a wide panel of possible environmental change by 2070 (SSP1 being now considered unrealistic and therefore, no longer recommended; (IUCN Standards and Petitions Committee, 2024).

### 2.4 Land use data

We used current data and future projections of land use at the global scale (resolution 0.05°, approximately 5 × 5 km; (Chen et al., 2020). To represent the overall absence of the species in forests and increase the realism of our projections, we performed a post-hoc filtering to remove all forested areas from the predictions of invasion risks. Provided the species is strongly associated (either negatively or positively) with a given habitat type, using a filter (or a mask) enables to account for habitat requirements without adding any habitat predictor to the models, therefore preserving statistical parsimony. We used current (2015) and future (2070) ensemble model projections (i.e. the mean projection across five GCMs) for the same scenarios as our climate data (RCP45-SSP2 and RCP85-SSP5; Fig. S1). We selected all predictors of temperate and tropical forest cover (expressed as percentages), i.e. temperate evergreen trees, tropical evergreen trees, temperate evergreen trees, tropical deciduous trees and temperate deciduous trees. We summed up all types of forest cover to produce a single map of percentage of tree cover for each projection (current, future SSP2 and future SSP5). To match the resolution of our projections (1 km), we resampled the tree cover maps with a bilinear interpolation. We set the predictions of invasion risks to zero where tree cover was over 95% (we only removed pixels which were nearly filled with forest because the species can be found in forest edges and glades, and because the vast majority of presence points were located in pixels with < 95% tree cover; Fig S2). To visualize the effect of future tree cover change (i.e. the difference between future and current tree cover), we used a mask of current and future tree cover, respectively, on our projections of future invasion risk, for each corresponding scenario.

### 2.5 Factors of spread

To account for the risk of introduction and spread of the species, we used anthropogenic predictors, which we assumed to be related to propagule pressure and spread. The species has been introduced accidentally through transportation of goods, notably horticultural exports and shipments of consumable goods (Gray, 2020; Moore, 2019; van den Burg et al., 2024), and spread within islands through transport of construction material (M.T. Ibouroi pers obs. 2024).

We used three types of drivers of propagule pressure and spread as proxies for factors of introduction and spread. We used distance to port and airports as a proxy for introduction risk. We obtained port data from the World Port Index (https://msi.nga.mil/Publications/WPI, accessed December 2024) and airport data from the OpenFlights Airport database (https://openflights.org/data.html, accessed December 2024). These data were computed as the Euclidean distance between all pixels and the closest port / airport. We used primary road density (which included highways and main roads) as an indicator of potential spread facilitation (Lanner et al., 2022), because the species is known to spread through vehicular transportation of goods. We used data from the Global Roads Inventory Project (GRIP4) dataset (Meijer et al., 2018). We included human population density as a proxy for both introduction and expansion risks, since the species has been intentionally released in the wild or escapes from pet owners (e.g. in Florida; Gray 2020). We obtained human population density data for the current period and all four future scenarios from Gao (2020) at a resolution of 1 × 1 km². We were unable to include future scenarios for distance to ports, airports and roads because such data are not available; thus, our projections assumed no change in these predictors.

### 2.6 Background selection

It is recommended to use a restricted background to mitigate the effect of sample bias, follow the hypotheses of past dispersal and account for the Equilibrium hypothesis (Acevedo et al., 2012; Dubos et al., 2022c; Vollering et al., 2019). In the case of invasive species that are still spreading, it is preferable to exclude areas that are potentially suitable to the species but have not yet been reached to avoid generating of pseudo-absences in these zones and prevent false negatives (i.e. suitable environments predicted as unsuitable). We defined the background as a rectangle of minimal surface that includes all presence points, which we extend by 10 % in each cardinal direction. This results in a map which includes Florida, central Africa, the Comoros Archipelago, Réunion Island and Mauritius (Fig. S3). After model training on this background extent, we projected the predictions of invasion risk on a global map, excluding cold temperate climates from both hemispheres (latitude > |40°|).

### 2.7 Pseudo-absence selection and sample bias correction

We generated 40,000 pseudo-absences with the spatSample function of the terra package (Hijmans, 2025). This number represents a fair trade-off between model reliability and computation time (Valavi et al., 2021). A former sensitivity analysis showed no difference with 10,000 pseudo-absences (Unpublished Master’s thesis, Calesse 2023). We generated three sets of pseudo-absence. We applied a sample bias correction technique consisting of a selection of pseudo-absence points that imitate the spatial bias of presence points (Phillips et al., 2009). To do so, we used a null geographic model (Hijmans, 2012), which is the most appropriate approach for species that are still spreading and have not reached equilibrium, because it downplays the areas that have not been invaded yet and therefore mitigates the effect of non-equilibrium. We used the null geographic model as a probability weight for pseudo-absence selection (Fig. S3).

### 2.8 Predictor selection

The selection of environmental predictors is an unresolved issue in SDMs (Leroy, 2023). To follow the latest methodological recommendations (Araújo et al., 2019; Gábor et al., 2019; Sillero et al., 2021; Sillero and Barbosa, 2021), we performed a predictor selection process in five steps: (1) removing predictors with low variability in the study region, therefore unlikely to explain the presence of the species (Araújo et al., 2019), (2) removing collinear predictors (Dormann et al., 2013), (3) keeping predictors with the clearest causal relationship with species presence and spread (Dubos et al., 2022d; Fourcade et al., 2018; Hui, 2022), (4) selecting predictors with the highest relative importance (Bellard et al., 2016; Thuiller et al., 2009), and (5) considering potential interactive effects (Gábor et al., 2019). (1) We discarded bio3 (isothermality) because the relationship with species presence was not straightforward (bio3 being an interaction between daily and yearly variation in temperature), and displayed little variability on small islands (hence, yielding little capacity to explain species presence). We kept the remaining 18 variables because they all may explain species distribution (mostly through direct effects of temperature and water balance on physiology, or indirect effects on ecosystem productivity and food availability (Dubos et al., 2018). (2) For each climate data source and for each species-specific background (details below), we selected one predictor variable per group of inter-correlated variables to avoid collinearity (Pearson’s r > 0.7; Fig. S4) using the removeCollinearity function of the virtualspecies R package (Leroy et al., 2016). (3) When mean values were collinear with extremes, we selected the variables representing extreme conditions (e.g., warmest/driest condition of a given period) because these are more likely to drive mortality and local extirpation and be causally related to the presence of the species (Maxwell et al., 2019; Mazzotti et al., 2016). (4) We excluded the predictors with the lowest relative importance (assessed by the GINI index of the RandomForest R package). Relative importance was assessed with centred and scaled predictors (Fig. S5). (5) The effect of temperature can differ according to the level of drought (Dubos et al., 2019). To consider potential interactive effects between temperature and precipitation while accounting for propagule pressure, we kept a final set of two predictors related to temperature and two related to precipitation (in addition to the selected factors of spread). The latter step implies that we may re-qualify predictors which were formerly excluded.

### 2.9 Modelling technique

We modelled and projected climatic niches using two algorithms evaluated as the best performing ones (Valavi et al., 2023, 2021), i.e. Random Forest down-sampled (hereafter RF down-sampled), i.e. RF parametrised to deal with a large number of background samples and few presence records (R package randomForest; Liaw and Weiner 2002; Prasad et al., 2006) and MaxEnt (R package Predicts; Hijmans, 2024; Phillips et al., 2009). We set RF down-sampled to run for 1000 bootstrap samples/trees, each iteration randomly samples the same number of pseudo-absences as the number of presences, with replacement permitted. The number of variables randomly sampled as candidates at each split (argument *ntry*) was set to 3. For MaxEnt, we allowed all available types of relationships (i.e. feature class) except threshold ones, because these are more likely to be driven by overfit. The regularization multiplier was set to 1, which is appropriate for large samples. After discarding poorly performing models (see *Model evaluation* section), we averaged all projections across replicates.

### 2.10 Model evaluation

We used block-cross validation approach which better assesses model transferability compared to random partitioning (Valavi et al., 2023, 2019). We spatially partitioned our data into four folds using blocks of 500 × 500 km (Fig. S6). We chose this size of block so that (1) each fold includes a variety of geographic regions in order to cover well the climate niche of the species, and (2) the number of presence points is balanced between blocks. We assessed model spatial transferability with the Boyce Index (CBI; Hirzel et al., 2006), because other discrimination metrics (e.g. AUC and TSS) require absence data and may be misleading with biased presence-only data (Dubos et al., 2022d; Leroy et al., 2018), especially in the case of IAS at non-equilibrium. An index of 1 suggests that suitability models predicted perfectly the presence points, a zero value suggests that models are no better than random and a negative value implies a counter prediction. We discarded all models with a Boyce index < 0.5. We present projections of the mean suitability obtained across all highly transferable models (total number of models: 4 block-cross validation runs × 3 pseudo-absence runs × 2 modelling techniques = 24; for future projections: 24 × 2 scenarios × 3 GCMs = 144 projections).

### 2.11 Quantifying the effect of environmental change

We quantified the projected change in environmental suitability at the presence points (a) of the native range and (b) of the three invaded regions separately. We extracted the predictions of current and future invasion risk to represent the effect of environmental change on current populations. To highlight the effect of projected deforestation, we extracted suitability scores from the forecast maps (future projections) which were filtered with current (2015) and future (2070) tree cover (see *Land use data* sub-section).

### 2.12 Quantifying climate niche overlap

We assessed the level of similarity between the climate conditions of the native and the non-native regions. We computed a Principal Component Analysis between the selected climate predictors (see *Predictor selection* subsection) of the native and non-native ranges following Broennimann et al. (2012). For each of the three invaded regions (Florida-Caribbean, Comoros and Mascarenes), we quantified niche overlap with Schoener’s D overlap index and visualized both climate niches (as represented by two principal components) using the *ecospat.plot.niche.dyn* function of the ecospat R package (Di Cola et al., 2017).

## 3. Results

We selected nine predictors (five climate variables and four factors of introduction and spread; Table 1). All models were highly transferable, with a mean Boyce index of 0.92 ± 0.09 SD. No model was discarded.

**Table 1.**
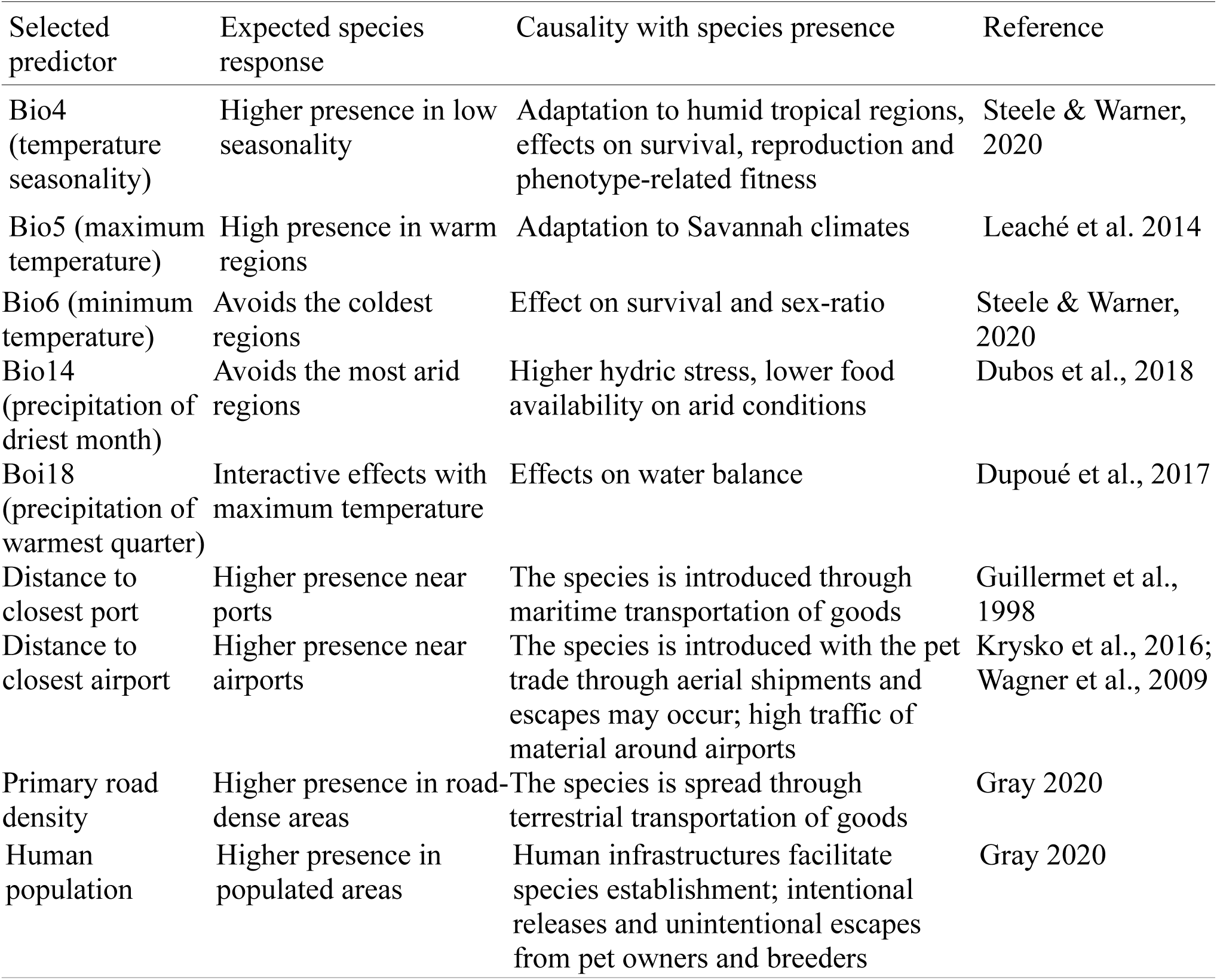
Selected environmental predictors of invasion risk for *Agama picticauda*, with expected responses and hypotheses of causal relationship with the presence of the species.

The species was absent from regions with a high temperature seasonality (Fig. S7). It established in warm regions (between 25 and 40°C of maximum summer temperature, Bio5), although the predicted values showed multiple modalities, reflecting differences between occupied regions. *Agama picticauda* is only present in regions with warm winters (> 13°C) and is absent from extremely arid regions (rainfall < 10 mm for the driest month, and < 100 mm for the warmest quarter; Fig. S7). Invasion risks are the highest near ports and airports (decreasing rapidly beyond 50 km), and are lowest in regions with low primary road densities and human populations. We identify several regions at high risk of invasion where there is no evidence of established populations yet, i.e. the Caribbean, East and Southeast Asia, the Brazilian-Argentine Industrial Core (corresponding to the MERCOSUR core economic region), Madagascar, Australia and Pacific Islands (Fig. 1).

**Fig. 1.**
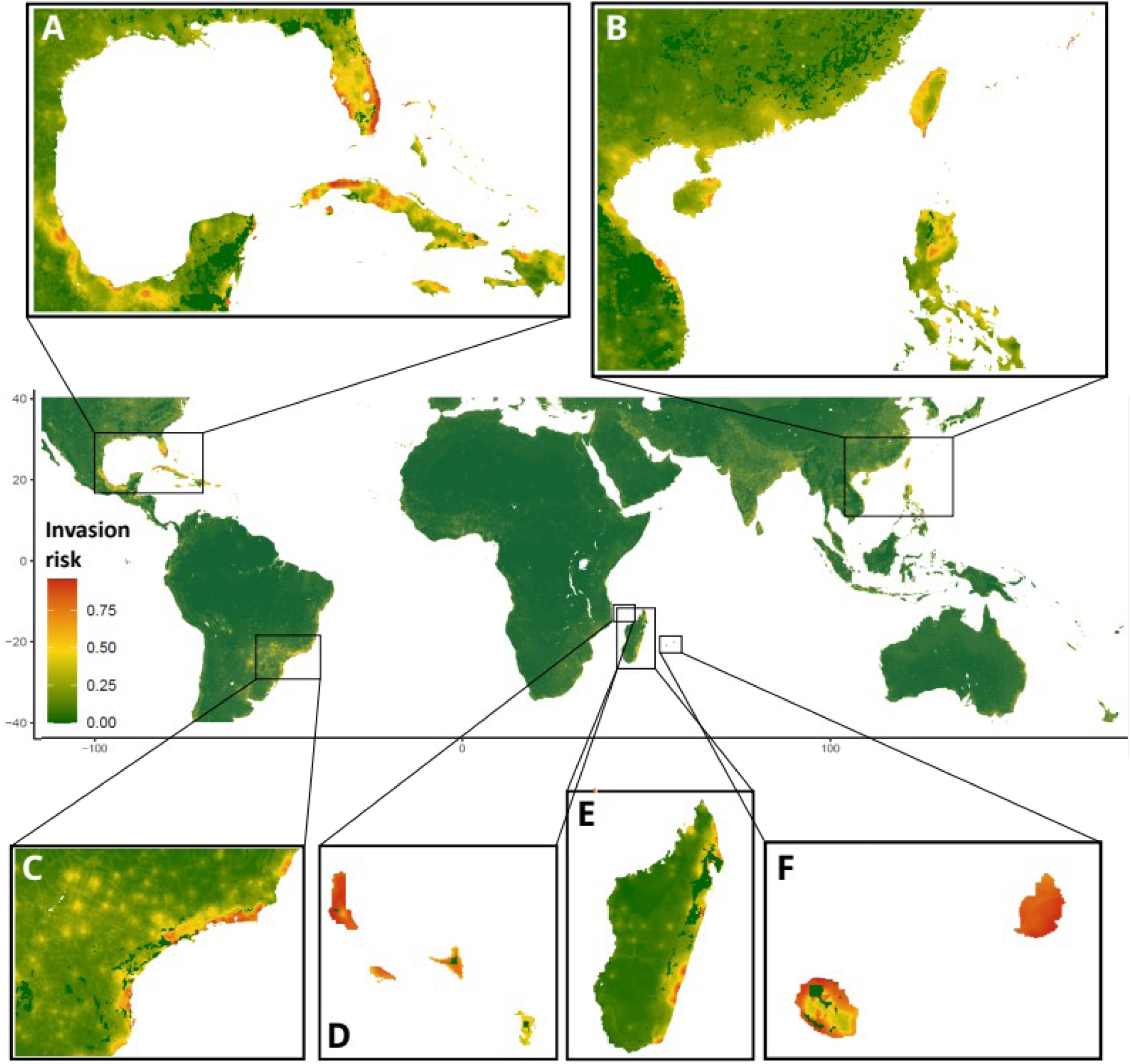
Global invasion risk of *Agama picticauda* under current conditions, projected from an ensemble model which accounts for climate, habitat and anthropogenic factors of introduction and spread. We highlight regions of high invasion risks (A: Florida-Carribean region; B: East-Southeast Asia; C: Brazilian-Argentine Industrial Core; D: Comoros Archipelago; E: Madagascar; F: Mascarene Archipelago). Red: high invasion risk; Green: low invasion risk. We show maps with presence points (Fig. S8) and the ensemble projections of individual algorithms (RandomForest down-sampled *versus* MaxEnt; Fig. S9) in the supporting information.

The combined effects of climate, land use and human population changes was variable through space (Fig. 2–5). Invasion risk may increase in human-populated areas, especially in specific locations of Central America, Mayotte, Réunion Island, Japan and South America. Future environmental change will decrease invasion risks in Florida and China overall (Fig. 2, 3). At the location of current populations, invasion risk will vary differently between regions (Fig. 4). The risk will increase in Comoros, decrease in Florida, and become more variable in the Mascarenes (Fig. 4). Tree cover will vary in either direction in the future and depend on the scenario. It will increase in the Florida-Carribean region and China (especially in the SSP5 scenario), which will contribute to the mitigation of *A. picticauda* propagation (Fig. 5, 6). Deforestation will create some new areas at risk, but mostly in areas that are not suitable. In regions with a high environmental suitability, deforestation will represent a small surface and are hard to perceive compared to climate and anthropogenic change (Fig. 5, 6).

**Fig. 2.**
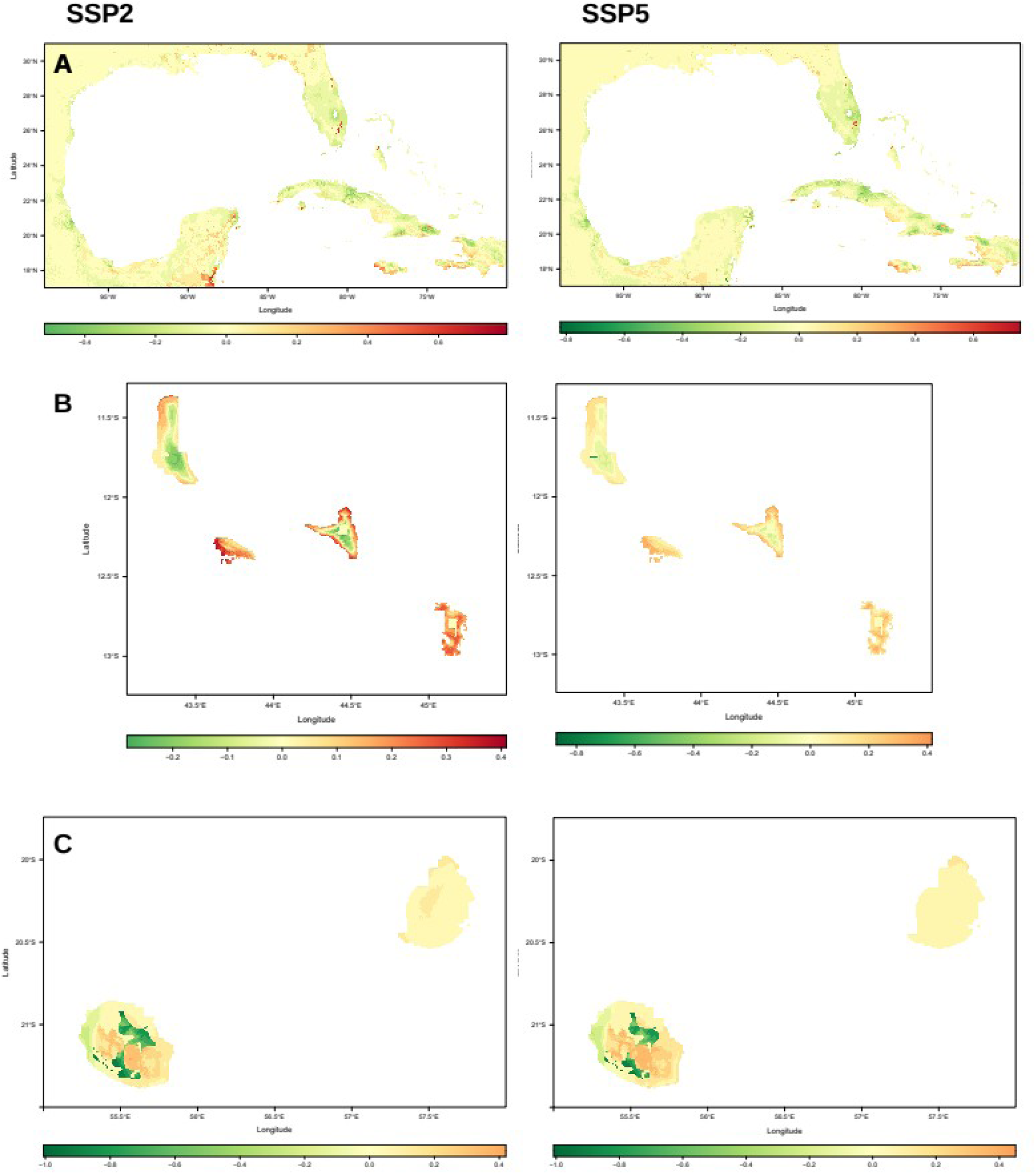
Effect of future environmental change (climate, tree cover and human population) on projections of invasion risk by *Agama picticauda* at three invaded regions and their surroundings (A: Florida-Carribean, B: Comoros Archipelago, C: Mascarene Archipelago). We show two scenarios (Left panels: SSP2, Right panels: SSP5) for the 2061-2080 period. Red: Increase in invasion risk; Green: Decrease.

**Fig. 3.**
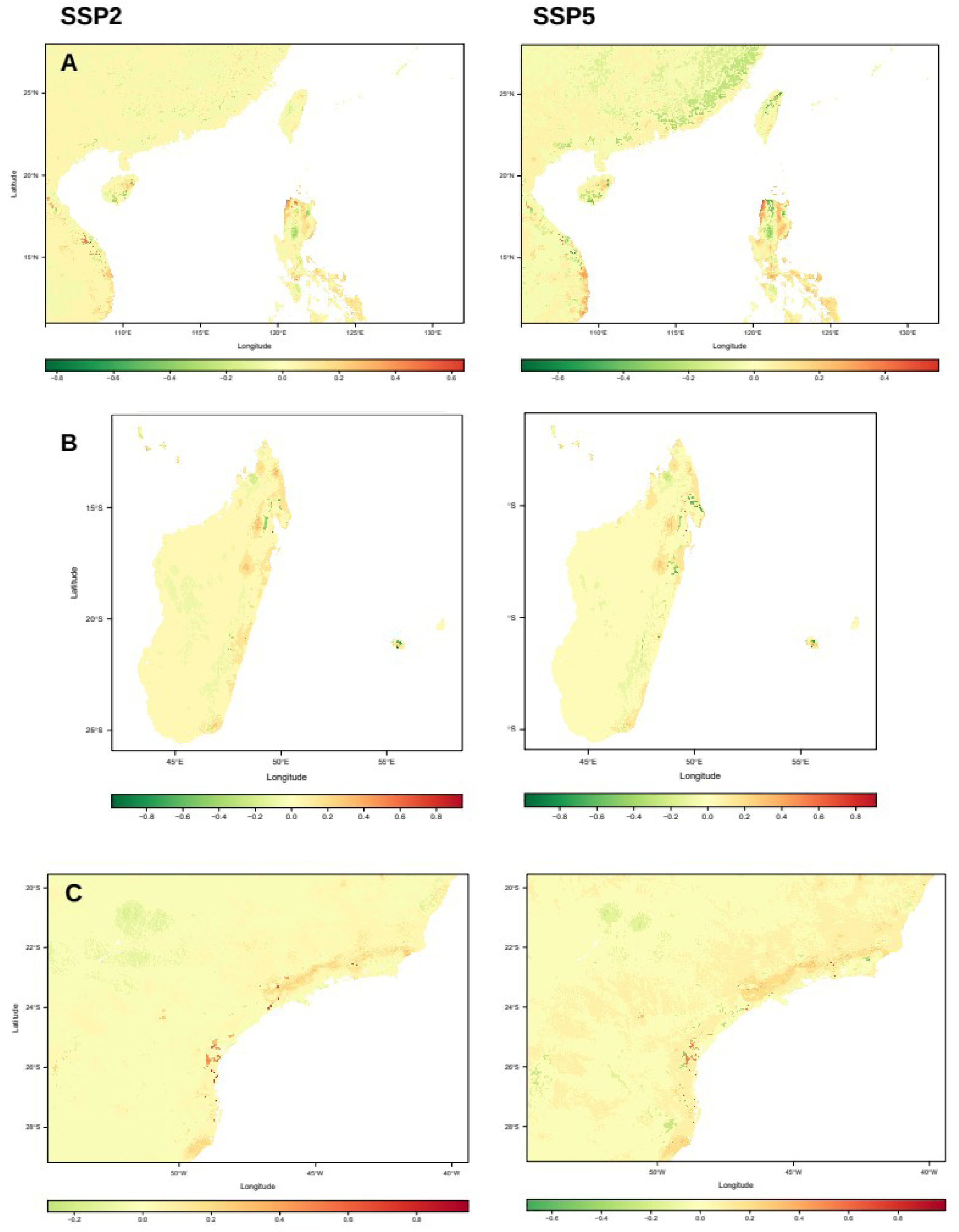
Effect of future environmental change (climate, tree cover and human population) on projections of invasion risk by *Agama picticauda* at three non-invaded regions (A: East-Southeast Asia, B: Madagascar, C: Brazilian-Argentine Industrial Core). We show two scenarios (Left panels: SSP2, Right panels: SSP5) for the 2061-2080 period. Red: Increase in invasion risk; Green: Decrease.

**Fig. 4.**
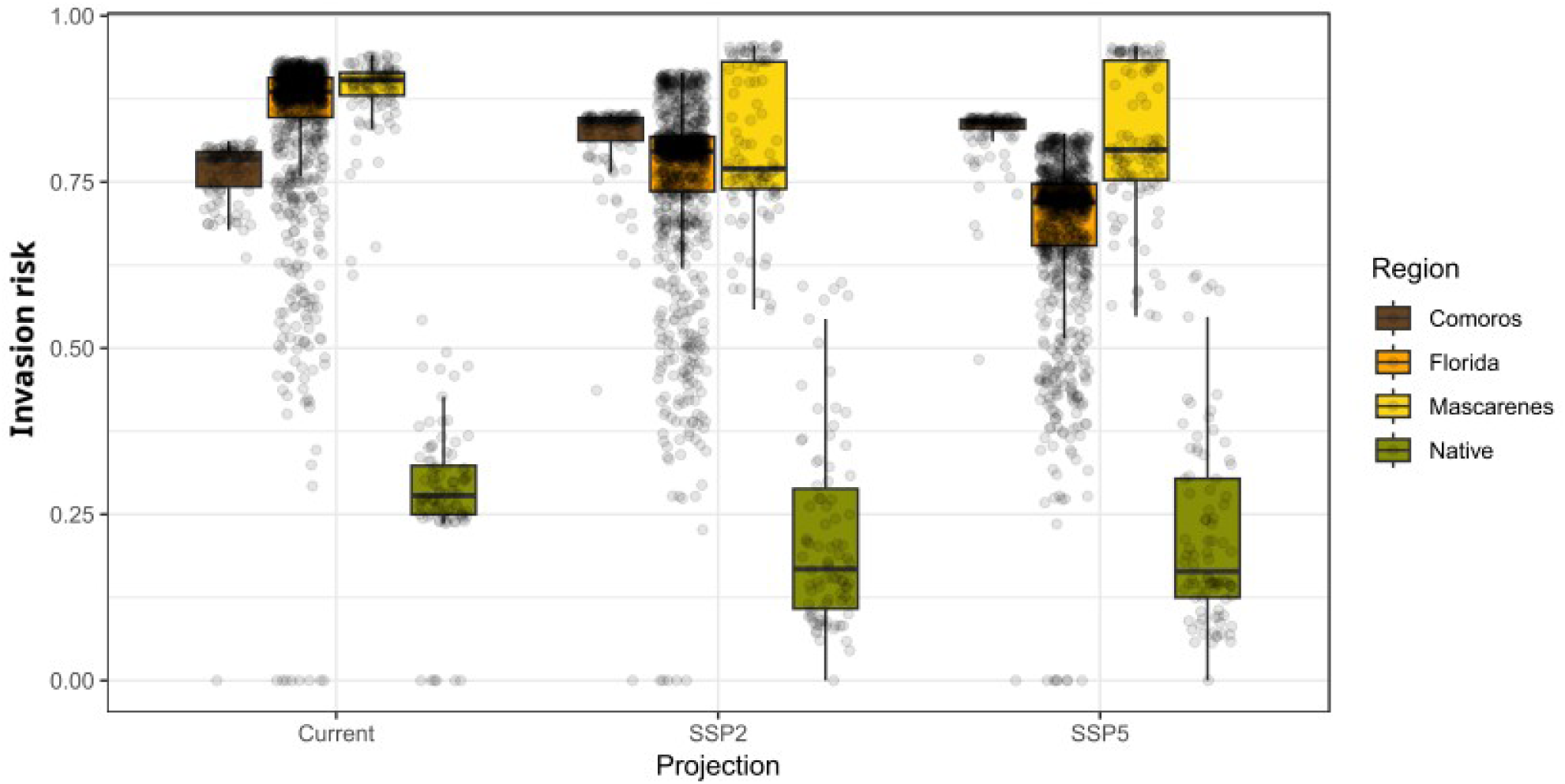
Predictions of current and future invasion risk (2070, two scenarios) by *Agama picticauda* at the current populations of the native (West-Central Africa) and the three invaded regions (the Comoros archipelago, Florida-Caribbean and the Mascarenes). Invasion risk was predicted from a combination of climate and habitat suitability, accounting for anthropogenic factors of spread. Boxes are composed of the first decile, the first quartile, the median, the third quartile and the ninth decile. Each point is a 1 km² occupied pixel.

**Fig. 5.**
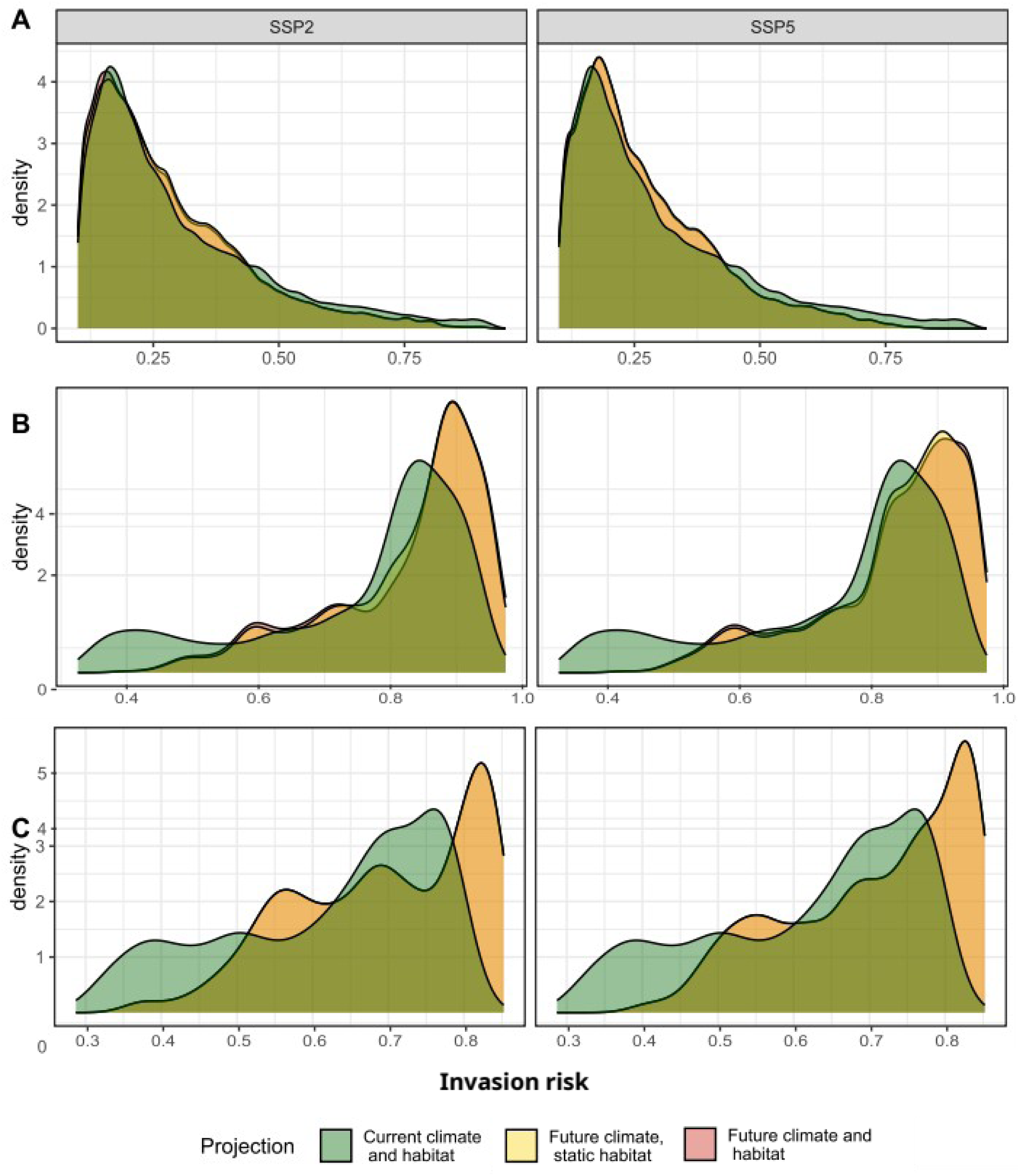
Distribution frequency of predicted invasion risk for three invaded regions (A: Florida-Caribbean; B: Mascarenes; C: Comoros), for current (green) and future conditions (2070, two scenario: SSP2 and SSP5), accounting for tree cover change (red) and assuming no change in tree cover (yellow).

**Fig. 6.**
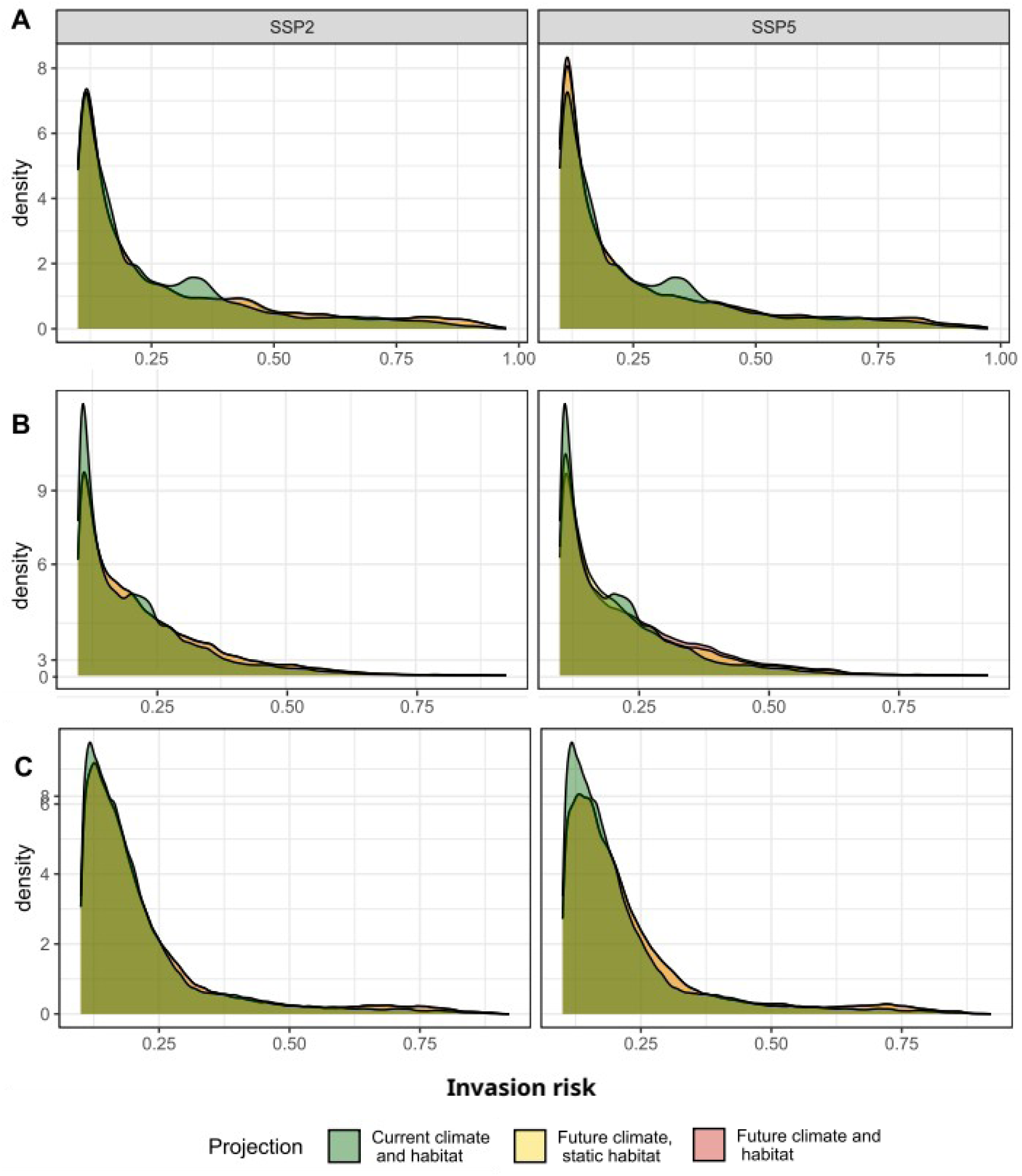
Distribution frequency of predicted invasion risk for three non-invaded regions (A: Madagascar; B: East-Southeast Asia; C: Brazilian-Argentine Industrial Core), for current (green) and future conditions (2070, two scenario: SSP2 and SSP5), accounting for tree cover change (red) and assuming no change in tree cover (yellow).

The niche overlap analysis revealed a high overlap between the native and the non-native regions, as shown by the blue areas of Figure 7. In non-native regions, the species is present in climates that are not represented in the native region (solid red areas), which provides evidence for niche expansion. In the Mascarenes and Comoros, some regions share the same conditions as found in the native range, but have not been invaded yet (indicated by the solid green areas; Fig. 7). This suggests that there is room for further spread in both regions. In Florida on the other hand, most of the conditions that are represented in the native regions are occupied, suggesting that the known realized niche is filled. However, the species has expanded its niche beyond the conditions available in the native range (light red) and might keep spreading. A significant part of the niche overlap between the native and invaded regions is related to anthropogenic factors of spread (i.e. human population and primary road density; Fig. S10).

**Fig. 7.**
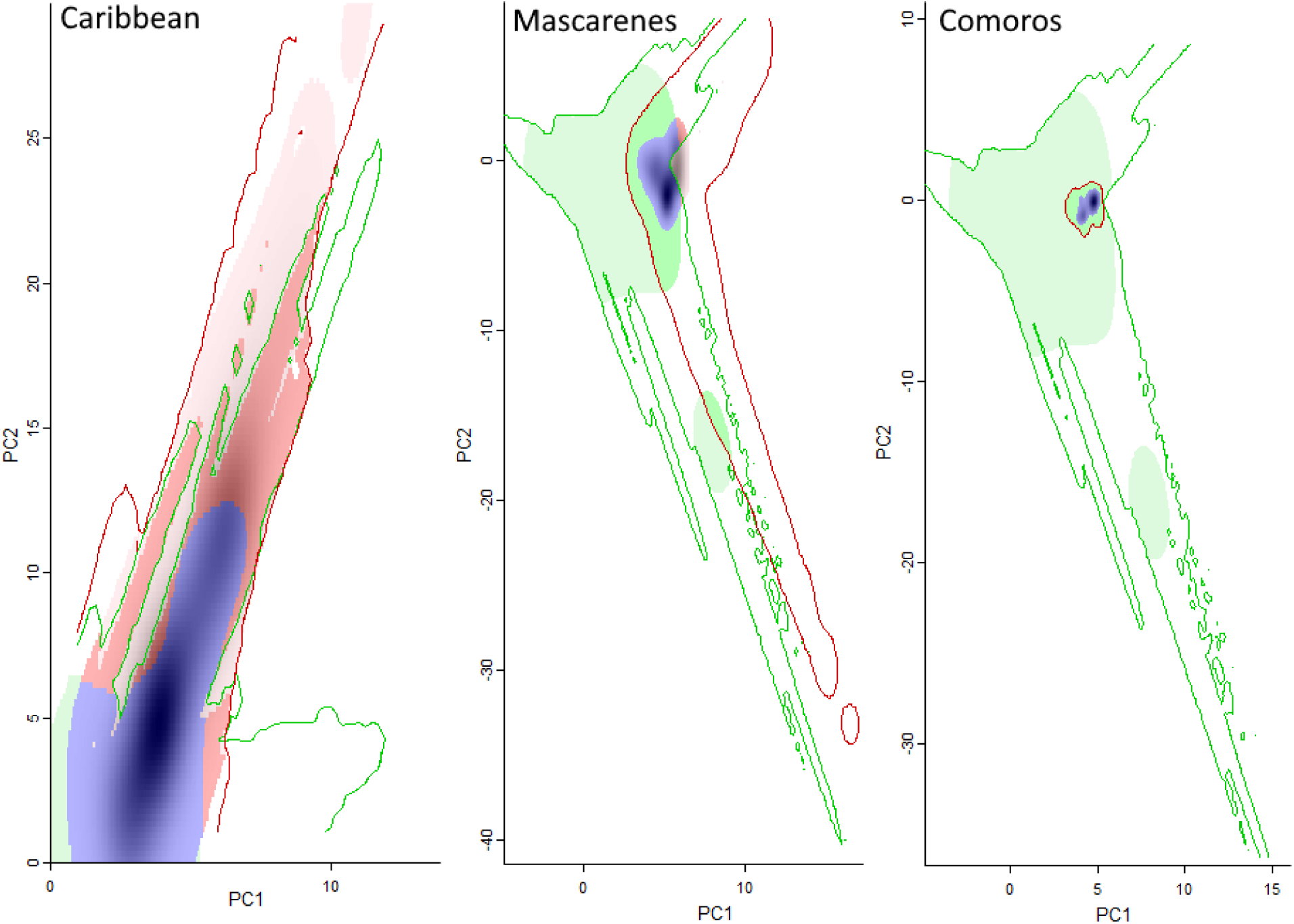
Realized climate niche of *Agama picticauda* obtained from Principal Component Analysis (five climate predictors and four anthropogenic factors of spread: bio4, bio5, bio6, bio14, bio18, distance to ports, distance to airports, primary road density and human population; see predictor contribution in figure S8). Each panel compares an invaded region with the native region (Florida-Caribbean, Mascarenes and Comoros). Green: climate niche of the native range (contour: environmental conditions of the native range; light green: occupied regions with conditions represented in the native region only; solid green: unoccupied regions of the non-native area with conditions that are represented in the native range); red: conditions of the non-native range (contour: environmental conditions of the non-native range, solid red: niche expansion under common conditions between the native and non-native regions, light red: niche expansion in novel conditions; blue: niche stability (i.e. overlapping conditions). The shades of black represent the density of occurrence points at a given set of principal component (PC) score.

## 4. Discussion

We predicted the current and future global invasion risks of *Agama picticauda*, accounting for climate, habitat and anthropogenic factors of introduction and spread. Global change will affect invasion risk, but in different direction and magnitude depending on the region. Our niche analysis indicated little room for further invasion in Florida and the risk should decrease in the future, apart from a few small surfaces predicted to be deforested. This contrasts with the Comoros, where the species may keep spreading, and where we predict an aggravation of the risk in the future. The risk will become more variable in the Mascarenes. We identify priority areas for proactive surveillance and highlight the need for region-specific management guidelines.

### 4.1 Drivers of invasion

#### 4.1.1 Anthropogenic factors

Factors of human-driven introduction and spread were the most influential predictors of the current distribution of *A. picticauda*, which is a common finding in studies of invasion risk (Bellard et al., 2016b; Lanner et al., 2022). The large amount of observation recorded from Florida, along with the important pet trade in this region may have largely contributed to the strong explanatory power of distance to ports and airports. The high density of presence points around airports suggests that many cases of introduction might result from escapes from the pet trade, as formerly observed for other marketed reptiles (e.g. *Phelsuma* spp.; Dubos et al., 2014). This can also be related to incidental introductions through the ornamental plant trade, as is the case for the green iguana (Villanueva et al., 2025). However, the first reports of *A. picticauda* in the three invaded regions were not in airports directly. In this specific case, the importance of anthropogenic factors was possibly due to intense vehicular movement of goods and vehicle traffic between highly populated urban areas and airports. The effect of human population (i.e. absence in unpopulated areas) likely reflects both the habitat use of the species and intentional and unintentional releases from the pet trade, and was mostly driven by Florida. *A. picticauda* is commonly found in human-modified habitats, where it can find shelter and food items (Mitchell et al., 2021). The increasing availability of human structures may benefit the species in the future and increase invasion risk in the modified regions (Gray, 2020). Invasion risks will increase at locations where human population is projected to increase, suggesting that anthropogenic factors may increase the chances of introduction and spread. This contrasts with the effect of climate change, which echoes another predictive study of invasion risks in the tropics (Dubos et al., 2025).

In the Comoros, the spread and colonization of new habitats by *A. picticauda* appear to be closely linked to urbanization and the expansion of built infrastructure (pers. obs. M.T.I., M.A.R.). One of the main vectors of dispersal is the inter-regional transport of construction materials, particularly sand, where the species lays its eggs. Additionally, porous concrete blocks used in construction often harbour eggs, juveniles, or even adult individuals, thus facilitating unintentional translocation. Although the species was initially introduced in the capital city of Moroni, it is now distributed across nearly all regions of the island. These findings suggest that human-mediated transport, especially through the construction sector, plays a key role in the dissemination of the species across the island landscape.

#### 4.1.2 Precipitation

The species was strongly influenced by summer precipitation, presumably due to water balance requirements during the warmest period. It is also possible that the avoidance of the most arid environments is related to indirect effects, through lower primary and secondary production, thus limiting food availability (Dubos et al., 2018). A decrease in summer precipitation might affect evaporative water loss in the hottest regions and impact negatively the persistence of the species.

#### 4.1.3 Temperature

*Agama picticauda* is currently distributed in regions with low temperature seasonality. This is consistent with the native region of the species, which is characterized by a tropical savannah climate (Peel et al., 2007). Fluctuating temperatures can affect sex-dependent survival and sex ratio (Steele and Warner, 2020). An increase in temperature seasonality may disrupt sex-dependent determination (towards more females; Steele and Warner, 2018) and affect population dynamics in the future. The establishment of the species in Florida may therefore seem surprising given the low temperatures occasionally occurring during winter. This suggests a high potential for behavioural or physiological plasticity or adaptation to new environments (Card et al., 2018; Fieldsend et al., 2021; Lapwong et al., 2021). Ecological release may also offer opportunities to colonize colder regions. This is the case for *Hemidactylus frenatus, Anolis sagrei, Leiocephalus carinatus* and *Phelsuma grandis*, which were able to spread in areas colder than their native range (Angetter et al., 2011; Claunch et al., 2021; Dubos et al., 2023a; Lapwong et al., 2021). Although our niche analysis indicated that the species already occupies all the environmental conditions that are represented in its native range, the width of its fundamental niche remains unknown, and *A. picticauda* may keep spreading towards novel conditions. This is supported by recent findings in new, colder locations (although establishment needs to be confirmed; Enge, 2024), and by the ability of reptiles to adapt or acclimate to colder conditions after experiencing cold snaps (Stroud et al., 2020).

Our models may have been limited by the fact that the species is still expanding its range. The data may not fully represent the entirety of the potential range of the species nor its niche extension. As a consequence, our projections of invasion risks may underestimate the potential spread of the species. We used the available data from multiple sources and recent sampling campaigns to inform models as much as possible, as recommended (Hui, 2022). Another limitation is model overfit. We observe some peaks in the predicted values of Random Forest (Fig. S7), but these do not reach extreme values, apart from bio5. The effect of overfit was mitigated by the combination of multiple predictors and with the ensemble modelling approach (combining Random Forest and MaxEnt). Besides, the spatial partitioning for model training and testing, along with high model performance indicated that models were highly transferable. The small amount of suitable environments at the global scale was presumably not caused by overfit, but rather by the limited surface represented by the combination of climate (tropical savanna) and anthropogenic infrastructure.

#### 4.1.4 Tree cover

We found that change in tree cover will not significantly increase invasion risk from a macroecological point of view. Most of the regions with projections of deforestation are located in regions of low environmental suitability. Tree cover is projected to increase overall in some regions (e.g. north Florida and eastern China), which might mitigate the spread. However, deforestation might create new areas available for establishment at the local scale in specific regions (e.g. In the Lesser Antilles; van den Burg et al., 2024). Accounting for tree cover may have increased the realism of our projections of suitable macroecological conditions. However, the resolution of our study (1 km²) may be limiting at the local scale, because *A. picticauda* is observed at the edge of large forests and in the surroundings of small tree patches, which cannot be perceived with our data. In addition, future projections may be uncertain, especially in regards to changes in local policies and stochastic events (e.g. in Comoros, where tree cover has been decimated across the archipelago in 2024 after a cyclone).

### 4.2 Niche analysis

We found that *A. picticauda* has colonized all the environmental conditions that are represented in the native region in the Florida-Caribbean region, including conditions of the native range where the species is absent. This supports that the species was constrained by biotic factors in the native region and suggests that niche expansion may be the result of ecological release (Fieldsend et al., 2021). Nonetheless, the species has extended its niche even beyond the conditions that are represented in Africa, suggesting either that the fundamental niche of the species is largely underestimated, or that the niche was enlarged through in situ adaptation (Fieldsend et al., 2021). In our specific case, a large proportion of the overlap is driven by road density and human population (Fig. S10), which emphasizes the importance of anthropogenic factors in explaining the current global distribution, especially in Florida. The situation contrasts in Comoros where we observe no niche expansion, and in the Mascarenes where niche expansion was low. The absence of niche expansion is presumably due to similar climates between western Africa and the archipelagos, along with the smaller size of the islands that limits potential variation in climate and habitats compared to continental regions. Nonetheless, we found a high potential for further spread in both regions. We showed that the Comoros may become completely invaded, since the available environmental conditions are all suitable to the species.

Although *A. picticauda* displays traits commonly associated with invasive species, such as ecological flexibility, territorial aggression and strong affinity for human-modified environments, there is hitherto little empirical evidence of direct negative ecological impacts. While *A. picticauda* has been identified as a potential threat to native herpetofauna, especially on islands with high endemism, there is currently no documented evidence linking its presence to confirmed population declines (van den Burg et al., 2024). This differs from arboreal IAS such as *Phelsuma laticauda* in Réunion Island (Deso et al., 2023), where there is greater niche overlap with endemic reptiles. However, the presence of *A. picticauda* on forest edges could affect arboreal species on small patches, which constitutes the majority of the habitat of some endemic species such as *P. inexpectata* (Dubos et al., 2022c). Native species would be better preserved at the core of larger forest patches, which supports the creation of large tree patches as a potential management solution.

### 4.3 Management recommendations

#### 4.3.1 Non-invaded regions

We identified priority for proactive surveillance at the global scale in regions that are still untouched by the species. Our models support that incidental escapes from the pet trade might be a major driver of invasion risk. We recommend to (1) increase biosecurity measures to promote early detection in ports and airports and (2) implement strong trade regulations to prevent further introductions, such as the banishment of *A. picticauda* from the pet trade in regions that are the most at risk (e.g. in the South American urban corridor, East and Southeast Asia). We recommend informing all stakeholders linked to the pet trade (including port managers, maritime transit, pet traders and pet owners) of the risk of escape and the subsequent ecological and economic hazards.

#### 4.3.2 Invaded regions

In Florida in particular, introductions are largely caused by deliberate releases from pet owners. This suggests that stronger regulations and public awareness may be a worthy investment to mitigating the risk in this region. Urban development is high in this region, which tends to create further opportunities for establishment (Clements et al., 2025). The species largely benefits from human structures and resources, either through the availability of shelters and food items, or through the low abundance of competitors (Mitchell et al., 2021). A key to slowing down the spread of the species is to promote the restoration of natural habitats. Such intervention may benefit both the native flora and fauna while reducing the amount of available space for *A. picticauda*.

In Comoros, the species is largely spread through transportation of building material. This calls for the need to integrate biosecurity into urban planning and increase surveillance between construction sites, logistic platforms and factories. Although representing a small surface at the global scale, deforestation may facilitate the establishment and spread of the species. We recommend the prevention of deforestation in the most sensitive areas (e.g. in the Caribbean, Mauritius and Mayotte). An increase in tree cover may be another potential way to mitigate the risk, provided native tree species are promoted. To mitigate the exposition of native arboreal species, the creation of large patches should be preferred over smaller patches. In regions that are exposed to cyclone activity such as the Comoros, extreme winds can deforest large areas and create colonization opportunities for *A. picticauda*. It is urgent to facilitate tree plantation in order to restore native habitats and prevent new establishments. Global change will increase the risk in the Comoros and Mascarene archipelagos, which stresses the need to account for future environmental suitability in habitat restoration plans. We recommend the development of monitoring programmes for *A. picticauda* populations, and the study of the response of the native vegetation to environmental change. Implementing these recommendations will require coordinated efforts among stakeholders, targeted awareness campaigns, and integration into local land-use planning and trade regulation frameworks.

## Supporting information

Supporting info Agama picticauda invasion risks

## Acknowledgements

We are grateful to all the staff of Nature Océan Indien for their help in data collection. Fieldwork in the Comoros was funded through a research support grant from the Rufford Foundation (grant no. 42957-1 awarded to Saoudati Maoulida).

## CRediT authorship contribution statement

**Nicolas Dubos**: Conceptualization, Data Curation, Formal analysis, Methodology, Investigation, Supervision, Writing – original draft. **Steven Calesse**: Data Curation, Formal analysis, Investigation. **Kathleen C. Webster**: Data Curation, Investigation, Writing – review and editing. **Chloé Bernet**: Supervision, Project administration, Funding acquisition, Writing – review and editing. **Gregory Deso**: Data Curation, Investigation, Writing – review and editing. **Thomas W. Fieldsend:** Investigation, Writing – review and editing. **Saoudati Maoulida**: Data curation, Writing – review and editing. **Soule Mounir**: Data curation, Writing – review and editing. **Xavier Porcel:** Data Curation, Investigation, Writing – review and editing. **Jean-Michel Probst:** Data curation, Investigation, Writing – review and editing. **Hindatou Saidou**: Data curation, Writing – review and editing. **Jérémie Souchet**: Supervision, Project administration, Investigation, Writing – review and editing. **Mohamed Thani Ibouroi**: Conceptualization, Data Curation, Investigation, Supervision, Writing – review and editing. **Markus A. Roesch**: Conceptualization, Data Curation, Methodology, Investigation, Supervision, Funding acquisition, Writing – review and editing.

