## Supporting info Agama picticauda invasion risks for "Diverging effects of global change on future invasion risks of *Agama picticauda* between invaded regions: same problem, different solutions"

By Dubos et al.

### Supporting information

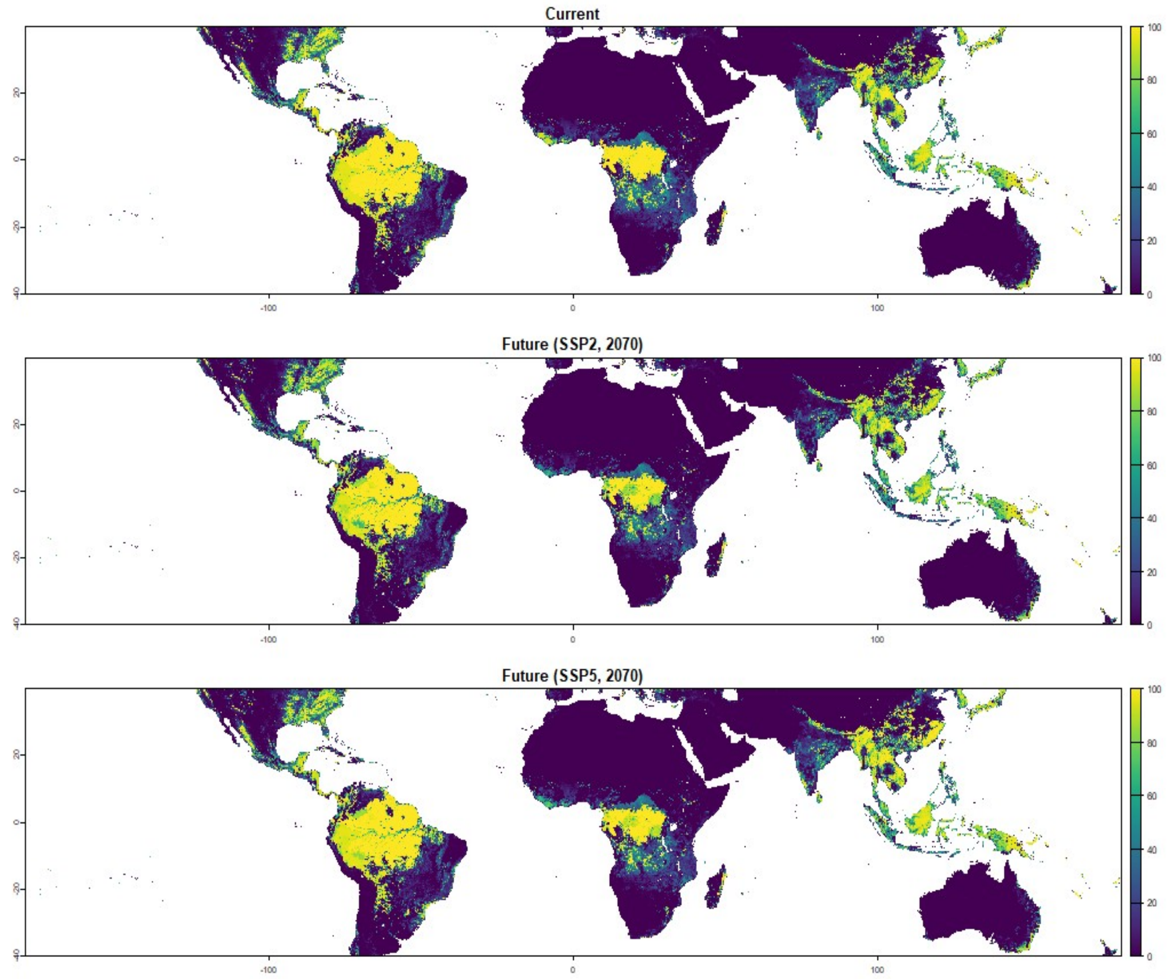


Fig. S1 Global tree cover for current (top) and future conditions (2070, two scenarios).

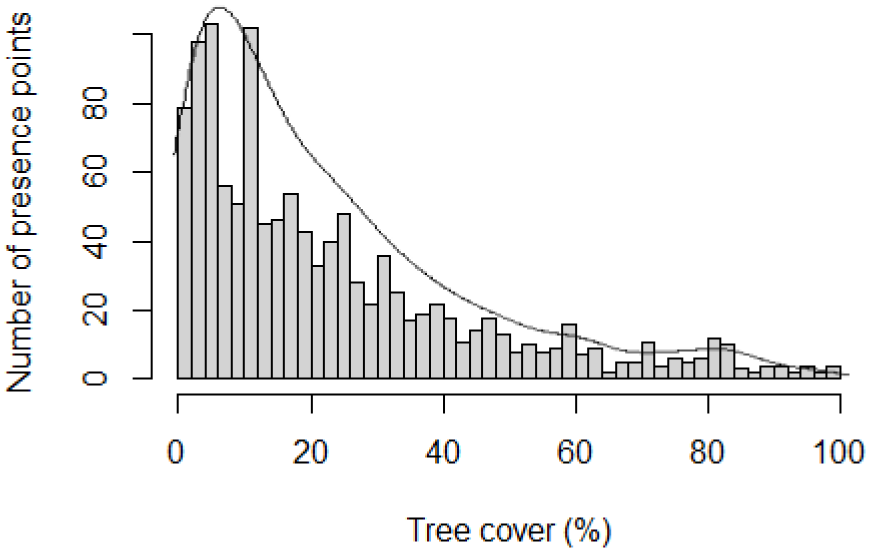


Fig. S2 Percent tree cover at the location of presence points for *Agama picticauda*.

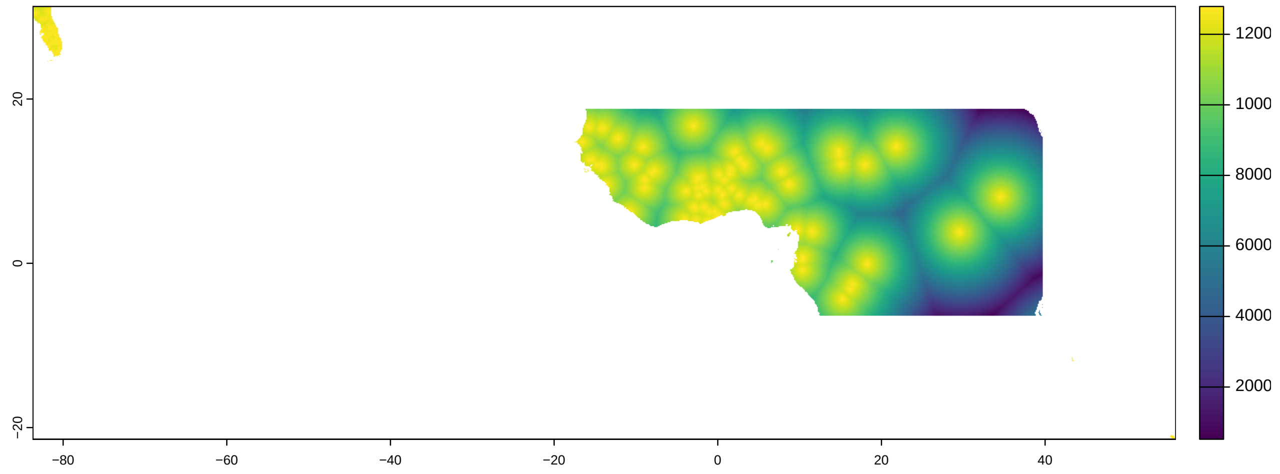


Fig. S3 Null geographic model used for sample bias correction. This map also shows the extent of the background (the area taken into account for model training). The geographic area includes central-western Africa, Florida, the Comoros, Reunion Island and Mauritius.

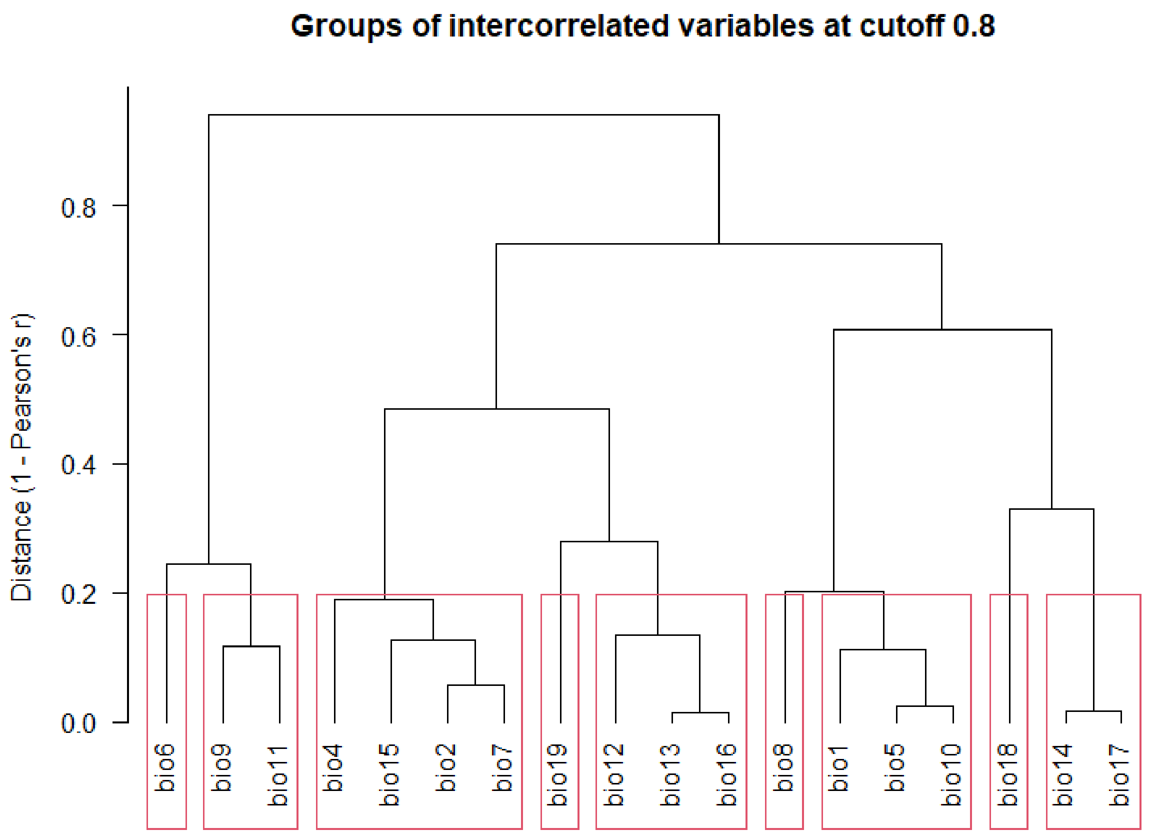


Fig S4 Predictor collinearity for the modelling background of *Agama picticauda*.

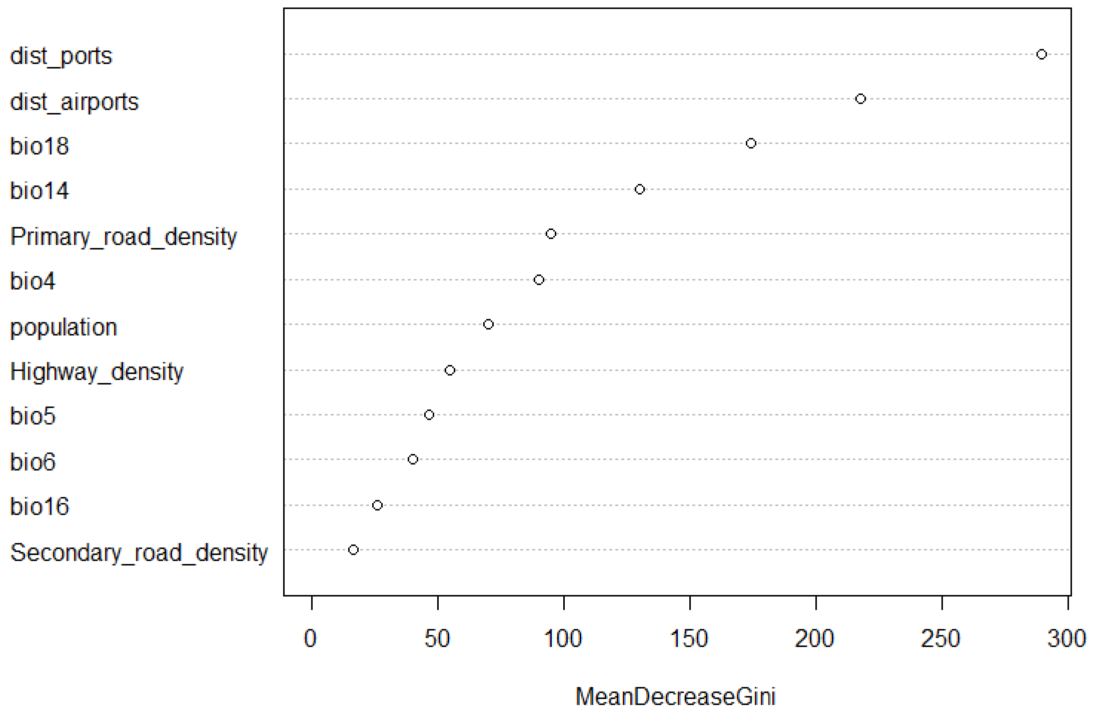


Fig. S5 Predictor relative importance for the presence of *Agama picticauda*.

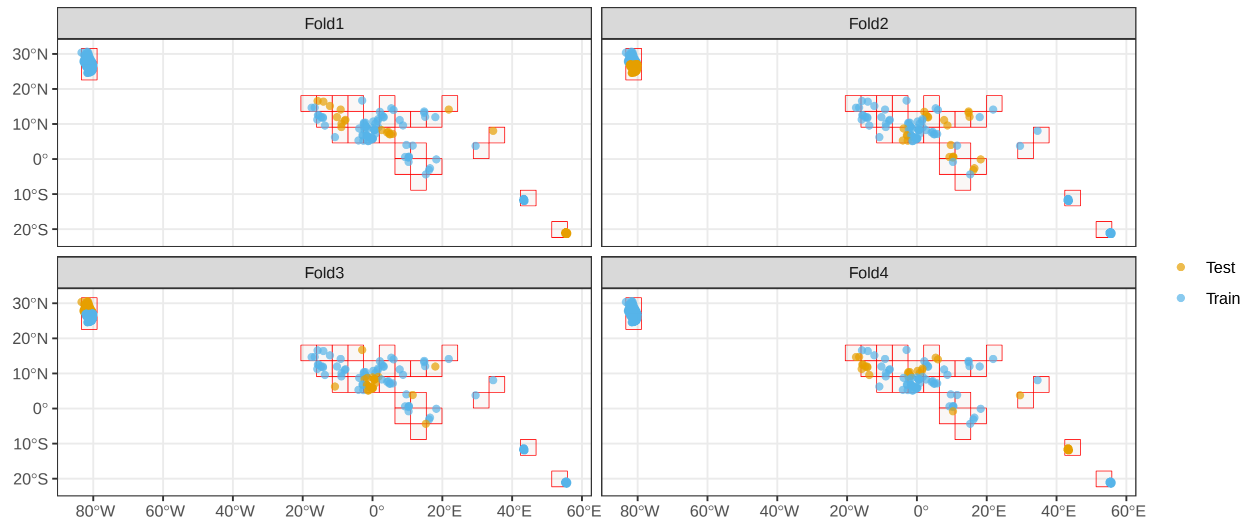


Fig. S6 Spatial partitioning for block-cross validation for invasion risk modelling of *Agama picticauda*.

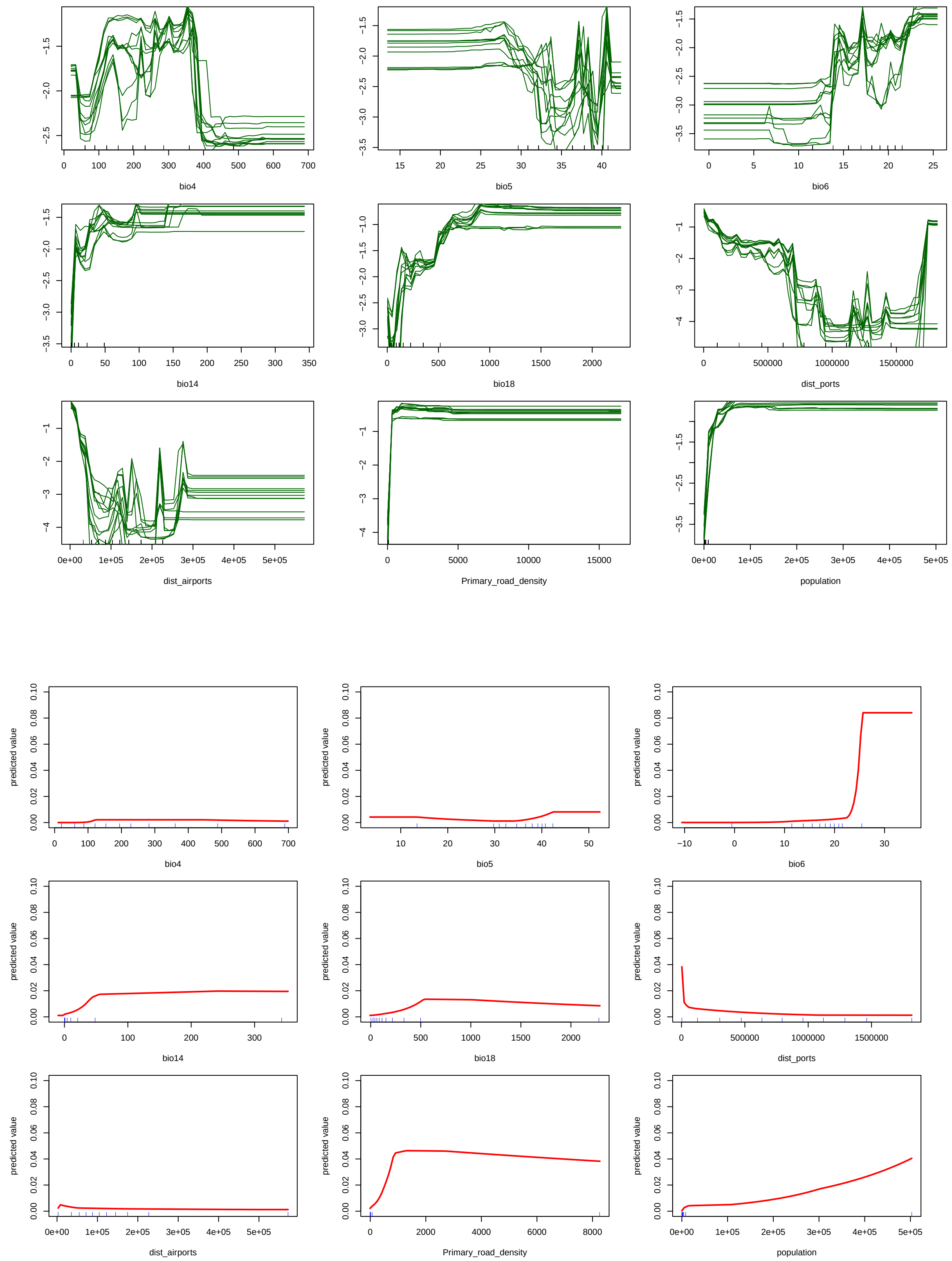


Fig. S7 Predicted response of *Agama picticauda* presence to environmental predictors (top / green: Random Forest down-sampled, bottom / red: MaxEnt). For Random Forest, each line represent one model iteration. For MaxEnt, we show one example of model iteration.

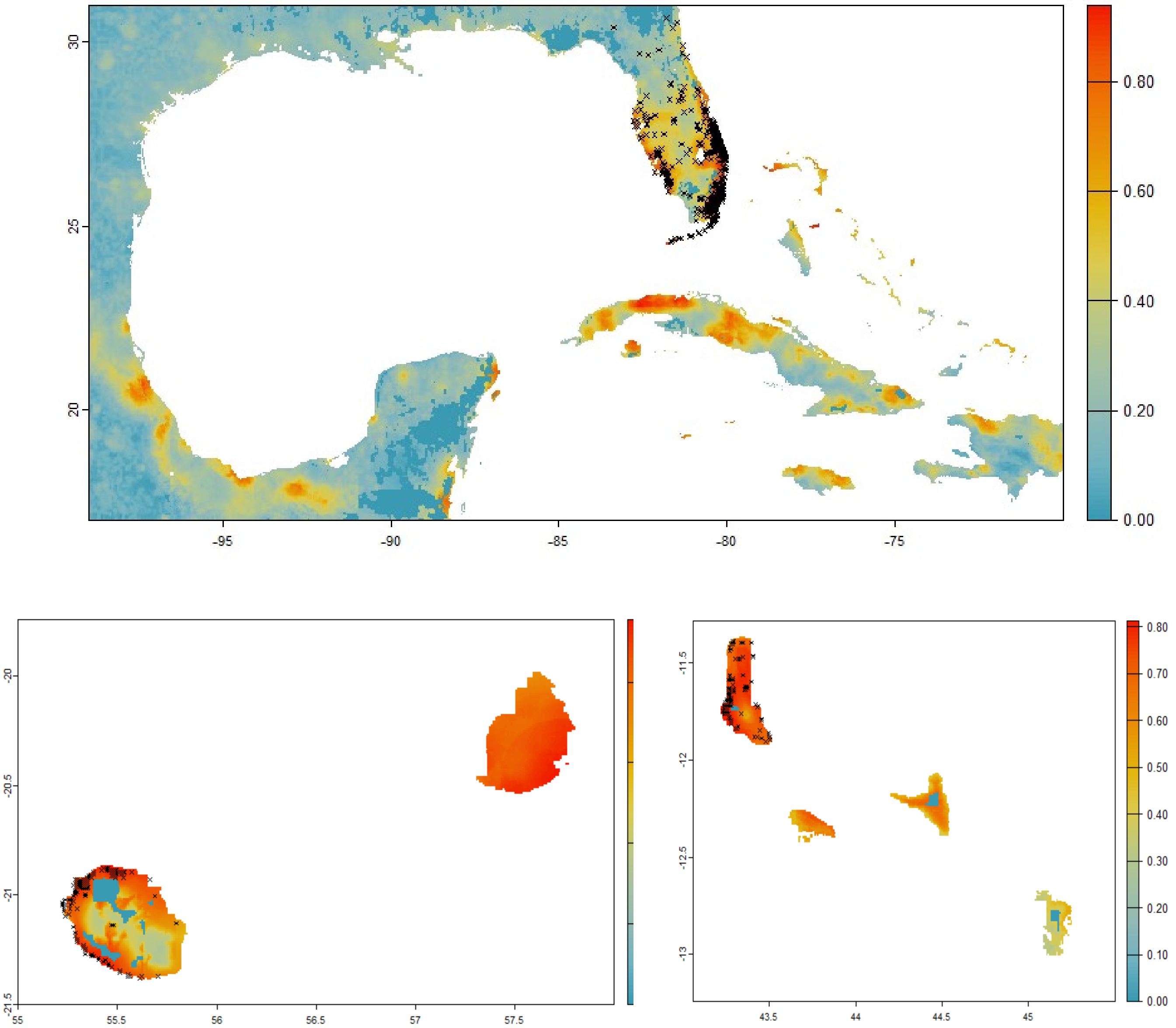


Fig. S8 Predictions of current invasion risk for three invaded regions (Florida-Carribean, Mascarenes and Comoros), with presence points (x).

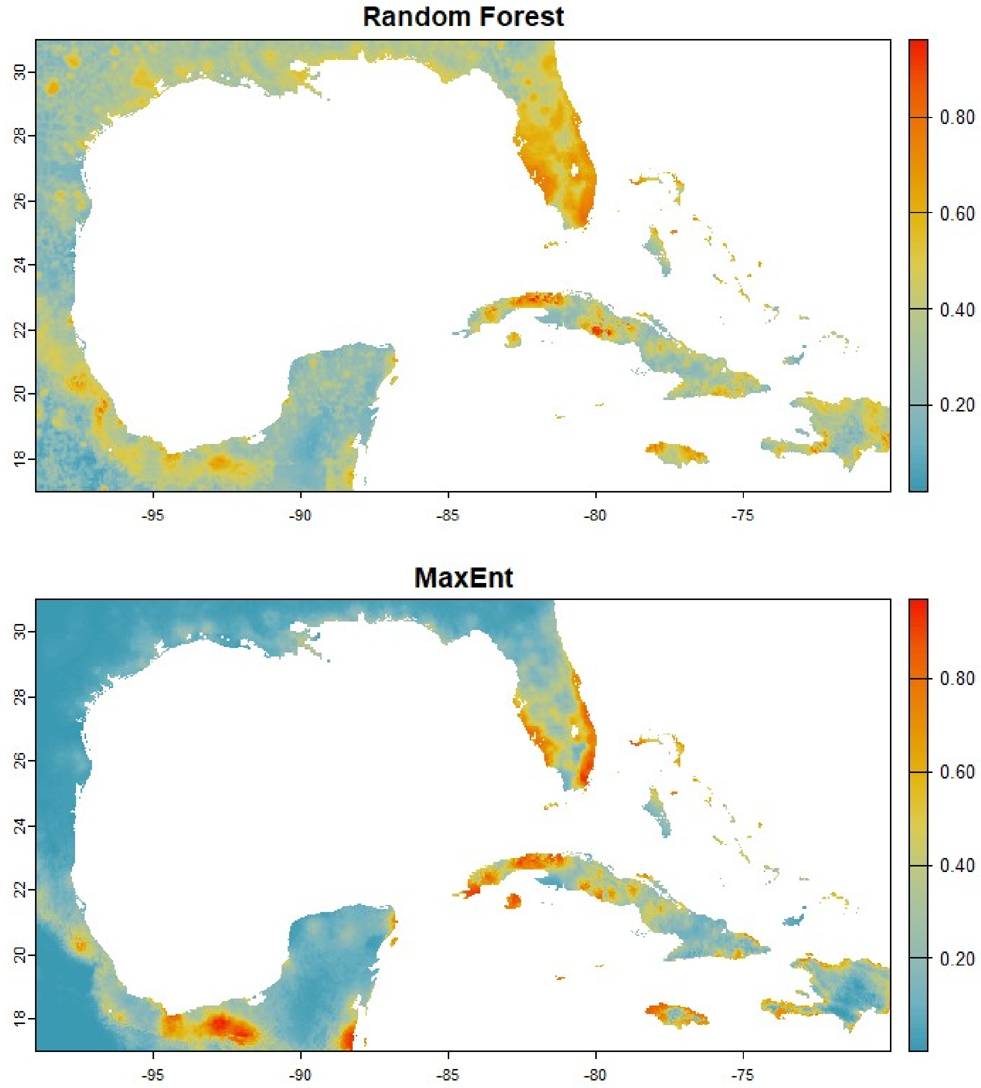


Fig. S9 Illustration of the effect of the choice of algorithm (Random Forest down-sampled *versus* MaxEnt), with the example of the Florida-Caribbean region. The two maps represent the average prediction across replicates of pseudo-absence and cross-validation runs.

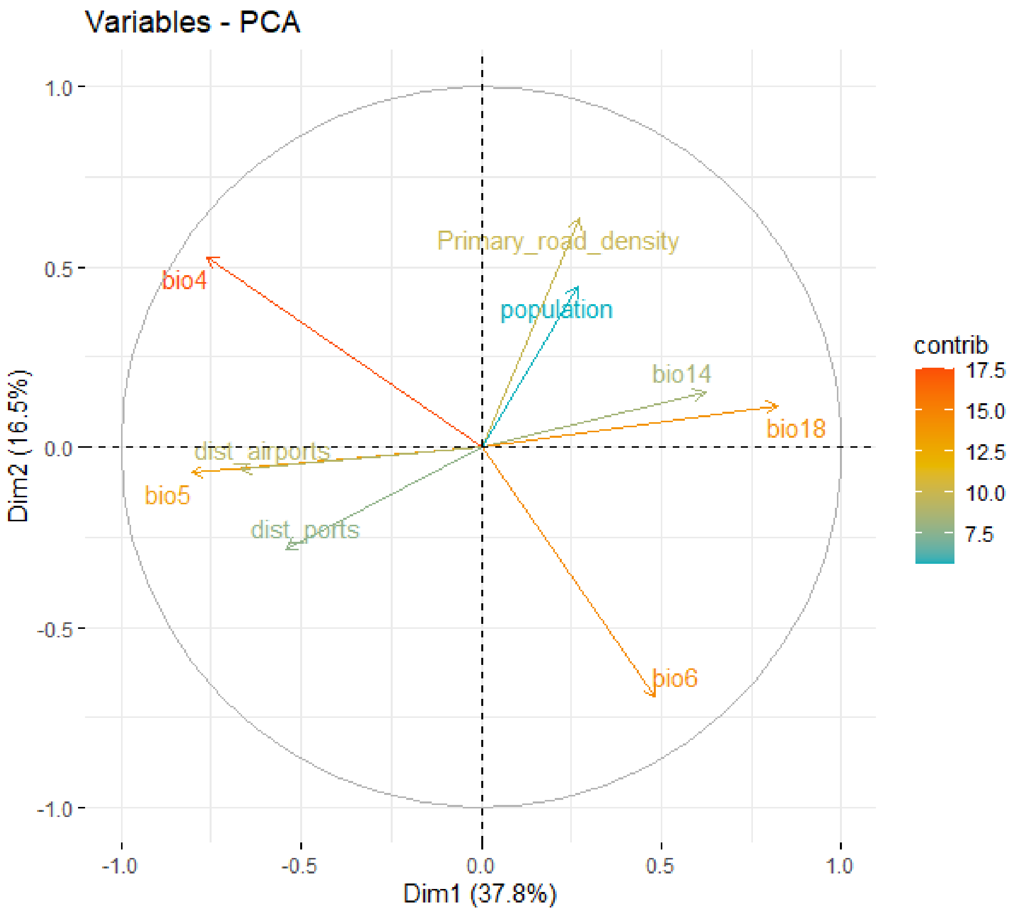


Fig. S10a Predictor contribution for the Principal Component Analysis of the realized niche of *Agama picticauda* – Region Caribbean.

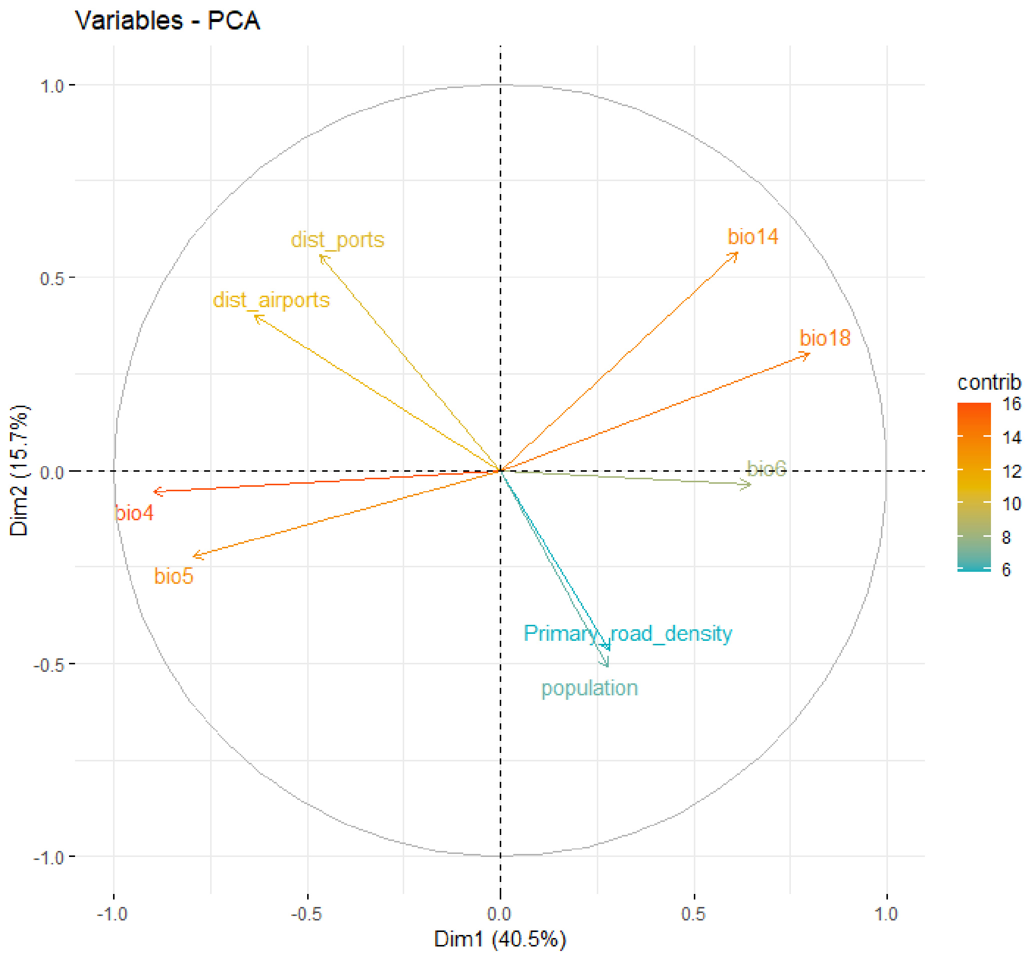


Fig. S10b Predictor contribution for the Principal Component Analysis of the realized niche of *Agama picticauda* – Region Comoros.

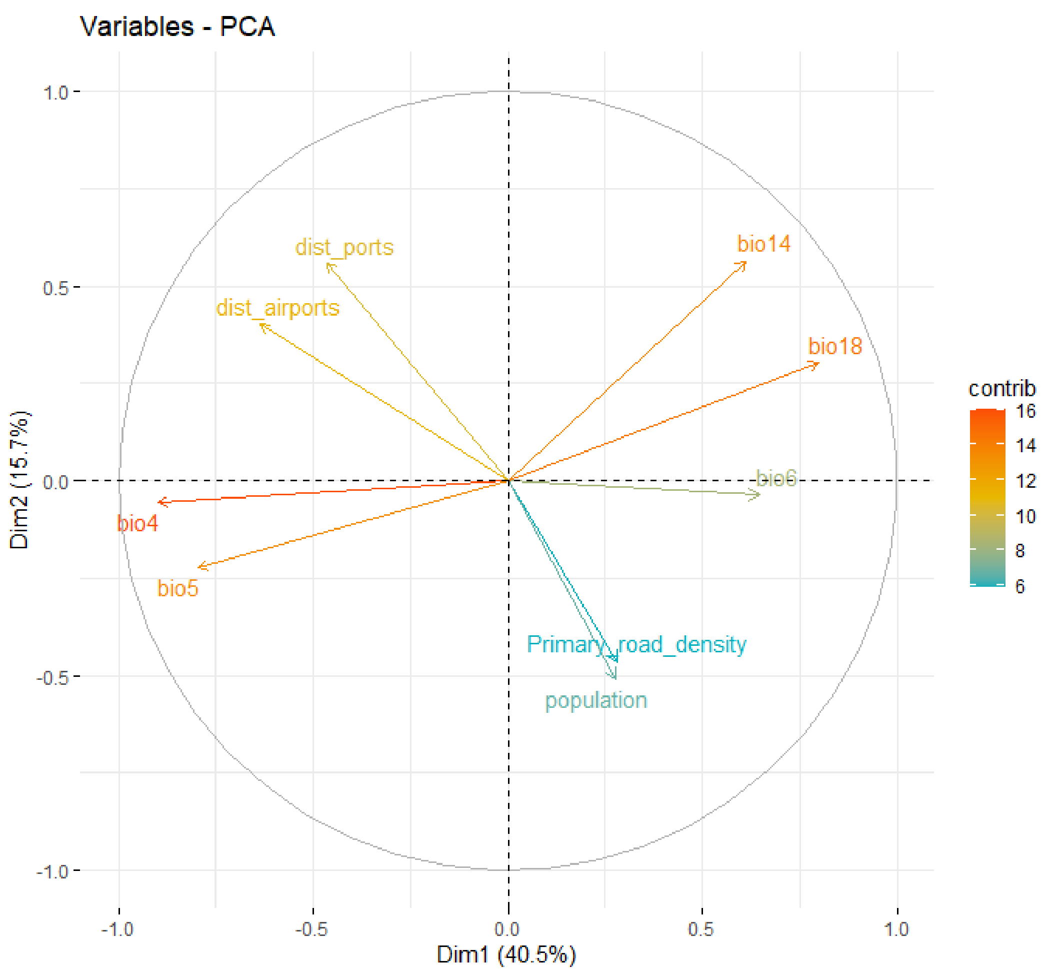


Fig. S10c Predictor contribution for the Principal Component Analysis of the realized niche of *Agama picticauda* – Region Mascarenes.
